# The essential function of the apical polar ring during the blood stage of *Plasmodium falciparum*

**DOI:** 10.64898/2026.01.08.698415

**Authors:** Pratima Gurung, Peter S. Back, Ilzat Ali, Gabrielle M. Hernandez, Jeffrey D. Dvorin

## Abstract

The apical polar ring (APR) is a defining cytoskeletal structure in apicomplexan parasites, critical for parasite morphology and host cell invasion. However, its molecular composition and function remain elusive in *Plasmodium falciparum*. Here, we identify and characterize PfAPR1 as an APR-resident protein. Conditional knockout of PfAPR1 reveals its essential role in asexual replication. Using iterative ultrastructure expansion microscopy (iU-ExM), we show that PfAPR1 predominantly localizes to the basal APR ring. Loss of PfAPR1 causes defects in daughter cell segmentation and subpellicular microtubules organization, while IMC formation and apical polarity are largely preserved. PfAPR1-KO parasites contact host red blood cells but fail to form a tight junction, resulting in a complete block in invasion. Using PfAPR1 as molecular bait, we identify additional APR proteins and delineate APR biogenesis with U-ExM. These findings define the molecular architecture and function of the APR in *P. falciparum,* highlighting it as a promising antimalarial target.

## Introduction

The unicellular apicomplexan parasite *Plasmodium falciparum* is the causative agent of the deadliest form of human malaria, claiming more than half a million lives globally^1^. *P. falciparum* replicates asexually in human red blood cells (RBCs) via schizogony and produces dozens of daughter cells (merozoites) per replicative cycle. This exponential parasite growth causes the symptoms of clinical malaria^2^. All apicomplexan parasites, including *Toxoplasma gondii* and *Cryptosporidium spp.*, *Plasmodium spp.* possess a characteristic structure called the apical complex^3–6^ for which they are named. This complex contains the apical polar ring (APR), secretory organelles (rhoptries and micronemes), and, in some apicomplexans, a conoid, along with distinct cytoskeletal structures^3,7^. Rhoptries and micronemes contain essential proteins required for motility, egress, host cell attachment, and invasion^8^. The conoid, a small cone-shaped structure containing tubulin-based filaments, is critical for host cell invasion in *T. gondii*. However, its presence in *Plasmodium spp.* is debated, with a conoid-like structure observed during the ookinete stage in mosquitoes^4,9,10^. The APR is a central component of the apical complex in all apicomplexans, identified by electron microscopy more than 50 years ago^3,4,11,12^. Together, the apical complex plays a crucial role in several essential process, including determination of parasite shape, egress out of the host cell, and invasion into a new host cell.

In *Plasmodium falciparum*, the APR is composed of three concentric rings with approximate diameters of 180 nm, 240 nm, and 370 nm. The two top apical rings are connected by a bridge structure consisting of up to 34 repeating units, while the third, more basal ring is the largest and thickest among the apical rings in merozoites^6^. This basal ring is connected to the parasite’s cytoskeleton, known as the inner membrane complex (IMC) and subpellicular microtubules (SPMTs)^6,10,13,14^. During schizogony, the IMC forms at the apical end of developing merozoites, which further expands and envelops them. It is presumed that the APR marks the apex of the IMC and has been hypothesized to serve as a microtubule-organizing center (MTOC) and/or anchoring site for the SPMTs in all apicomplexans^4,6,9,14–18^. Recent studies challenge the putative MTOC role of the APR and instead favor a model where the centrosome-associated MTOC initiates microtubule formation with later hand off of the SPMTs to the APR^19^. Notably, *P. falciparum* exhibits variability in the number of SPMTs across different invasive stages, including merozoites (1-4)^10,20^, sporozoites (∼16)^21^, and ookinetes (∼60)^9,22^.The APR and its associated cytoskeletal network are crucial for maintaining parasite shape and facilitating key processes such as cell division and host cell invasion. Despite its prominent presence as densely stained rings at the apical end of its invasive stages^6,10,20,23^, the molecular components and function of the *P. falciparum* APR remains largely unexplored.

Among apicomplexans, the APR has been extensively studied in *T. gondii*^4,9,18,24–28^. Similar to *P. falciparum*, *T. gondii* has three apical polar rings with similar structural organization. However, cryo-electron tomography studies show that its apical rings have more repeating units (∼ 40-45) compared to those in *P. falciparum* merozoites^6^. Interestingly, only four out of eight known APR components of *T. gondii* (APR3/TGME49_208340, APR4/TGME49_219500, APR2/TGME49_227000, and APR7/TGME49_320030) have orthologs in *Plasmodium spp.*^4,26^. In the rodent malaria parasite, *P. berghei*, the ortholog of TgAPR3, PbAPR1/PBANKA_0907700, and of TgAPR7, PbAPR2/PBANKA_1334800), have been localized as a ring at the apical end of ookinetes^4^. Additionally, PbARA1/PBANKA_1410950, an apical ring associated protein has been identified to localize to the APR of *P. berghei* ookinetes by immunoelectron microcopy^29^. This protein is unique to *Plasmodium spp.* without clear orthologs in *T. gondii* or other apicomplexans, suggesting species-specific adaptations in APR composition that vary among Apicomplexans.

Recently, PyAPR2 and two associated proteins, PyAPRp2 and PyAPRp4, were characterized as APR resident protein in the rodent malaria parasite, *P. yoelii*^16^. While PyAPR2 is dispensable for asexual development, its depletion causes severe fitness defects during ookinete development. The loss of PyAPR2 does not affect the formation of the IMC or SPMT. However, it impairs the formation of the APR by disrupting the radiating spines, creating an abnormal gap between the APR and the IMC in ookinetes. Defective apical anchorage of the SPMTs is also observed, ultimately preventing parasite transmission in mosquitoes. Additionally, this study reported eight PyAPR2 interacting proteins that localize to the APR in these rodent malaria parasites. Notably, depletion of PyAPRp2 or PyAPRp4 resulted in similar phenotype to PyAPR2 depletion in *P. yoelii*^16^. However, no APR proteins have been identified in *P. falciparum* and the biological function of the APR in human malaria causing parasites remains unknown.

In this study, we characterize PfAPR1 (PF3D7_1141300) as a *bona fide* APR-resident protein in *P. falciparum*. PfAPR1 forms a ring-like structure at the apex of the parasite’s apical complex. Functional studies reveal that PfAPR1 is essential for asexual replication, with a complete loss of host cell invasion. Using PfAPR1 as a molecular handle, we identify three additional components of the APR and shed light on the biogenesis process underlying this critical APR structure.

## Results

### PfAPR1 is an APR resident protein in *P. falciparum*

We leveraged the catalog of APR-localized proteins previously identified in the rodent malaria parasites, *P. berghei* and *P. yoelii*, to identify conserved yet uncharacterized APR components in *P. falciparum*^4,16,29^. We found PfAPR1 as a potential candidate following a BLAST search against the *P. falciparum* genome via PlasmoDB^30^. PfAPR1 exhibits 44.9 % sequence identity and 63.9 % sequence similarity with PbAPR1(PBANKA_0907700) **(Extended Data Fig. S1)**. PfAPR1 is ∼111 kDa (1,012 amino acids) containing pleckstrin homology (PH)-like domain at residues 876-977. It is predicted to be essential for asexual replication based on transposon insertion screens in *P. falciparum*^31^ and *P. knowlesi*^32,33^. To investigate the role of PfAPR1, we generated an inducible knockout (iKO) parasite strain (referred to as PfAPR1^DiCre^) in the NF54^DiCre(*pfs47*)^ background, which expresses split dimerizable Cre recombinase^34^. We replaced the native PfAPR1 gene locus with a loxP-flanked, codon-altered PfAPR1 tagged at the C-terminus 3 copies of the hemagglutinin (HA) epitope **(Extended Data Fig. S2a)**. The PfAPR1^DiCre^ parasite strain was verified by PCR, whole-genome sequencing, and western blot **(Extended Data Fig. S2b and S2c)**.

Immunofluorescence assays (IFAs) across the asexual blood stages revealed PfAPR1 is undetectable prior to the schizont stage. During schizogony, PfAPR1 appear as a distinct ring-like structure at the apical end of developing and mature merozoites **(Fig. 1a and Extended Data Fig. S2d)**. To more precisely determine the timing of PfAPR1 expression, we co-stained with antibodies against PfGAP45, an IMC-associated protein that localizes to the newly forming IMC during the early schizont stage (∼5 nuclei, 40 hours post invasion/hpi) and later surrounds mature merozoites^35,36^. PfAPR1 expression coincided temporally with PfGAP45 and appears at the apex of PfGAP45 in merozoites **(Fig. 1a)**. Notably, PfAPR1 also localizes more apically than the micronemal apical membrane antigen 1 (PfAMA1)^37^. This suggests that PfAPR1 resides at the extreme apical tip of the merozoites, possibly at the APR **(Fig. 1b)**.

**Fig. 1:**
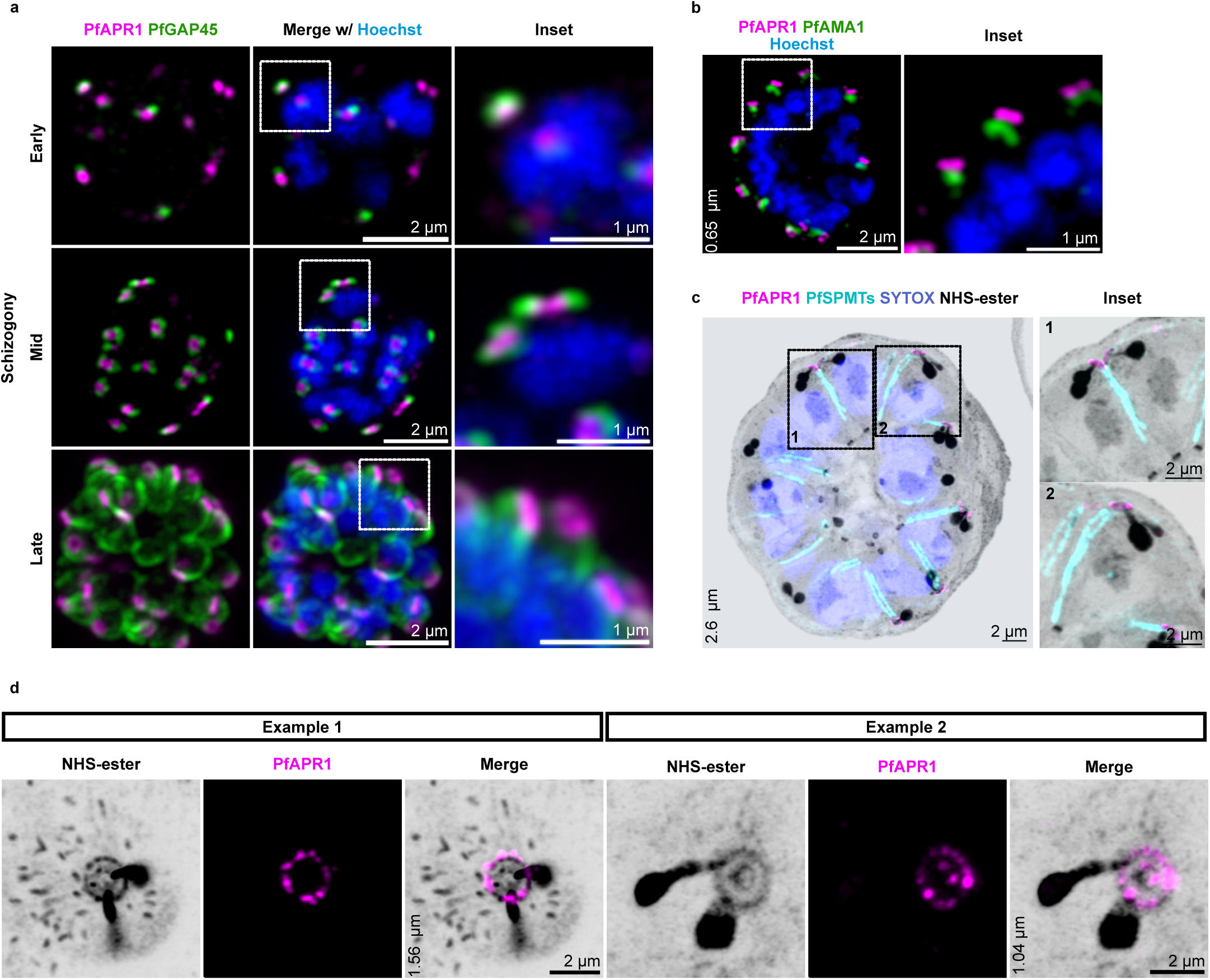
PfAPR1 is an APR-resident protein in asexual blood stages. Synchronized PfAPR1^DiCre^ parasites were analyzed during schizogony by IFA and U-ExM. **a,** Smear-based IFA images (maximum intensity projection) show HA-tagged PfAPR1 (ɑ-HA, magenta) as a distinct ring-like structure at the apical end of developing merozoites (early to late schizogony). PfAPR1 is localized at the apex of the IMC-associated marker PfGAP45 (green), which also serves as a marker for schizont progression. **b,** Batch IFA demonstrates that PfAPR1 (ɑ-HA, magenta) is positioned apical to the microneme marker, PfAMA1 (green) at the extreme tip of developing merozoites. **c,** U-ExM images of ML10-arrested late schizonts reveal that minus ends of SPMTs (ɑ-tubulin, cyan) are located beneath both PfAPR1 (ɑ-HA, magenta) and the NHS ester-stained APR, indicating spatial proximity to apical end in PfAPR1^DiCre^ parasites. **d,** Iterative U-ExM top view images show two representative examples: PfAPR1 predominately localizes to the basal APR ring is occasionally observed in upper APR rings as well. Regions within dashed rectangles are shown as zoomed inset. Results are representative of three independent biological replicates. Hoechst (blue) and SYTOX (blue) are used to stain nuclei in IFA and U-ExM, respectively. NHS-Ester (grey) is used as a general protein stain in U-ExM. Scale bars: 2 µm. For U-ExM images, z-axis thickness for maximum intensity projection is indicated in µm.

Using ultrastructural-expansion microscopy (U-ExM) with NHS-ester staining to label protein-dense APR^35,38^, we found that PfAPR1 forms a discrete bead-like ring structure overlapping the NHS-ester-dense APR. The minus ends of SPMTs are positioned beneath both PfAPR1 and the NHS-ester-stained APR, with their apical ends coinciding with PfAPR1 **(Fig. 1c)**. Previous ultrastructural studies have described the APR in *P. falciparum* as comprising three concentric rings^6^, whereas our U-ExM approach only resolved the APR as a single ring. To improve resolution, we applied iterative U-ExM (iU-ExM) with ∼15-fold expansion, which allowed us to distinguish two concentric APR rings: a large basal ring and a more apical, dense ring that likely represents the two top rings merged together, using NHS-ester. Notably, PfAPR1 predominantly localized to the largest basal ring and was occasionally observed in the apical ring (**Fig. 1d)**. Together, these findings establish PfAPR1 as a *bona fide* resident of the APR in *P. falciparum*.

### PfAPR1 is essential for asexual parasite replication

To investigate the role of PfAPR1 during asexual replication, we induced gene excision in PfAPR^DiCre^ parasites with rapamycin and confirmed excision by PCR and western blot **(Extended Data Fig. S2b and S2c).** PfAPR1-knockout (PfAPR1-KO) parasites displayed a severe growth defect, with 98 ± 0.6 % reduction in replication, within a single replication cycle compared to wild-type (WT) parasites **(Fig. 2a)**. Hemacolor-stained smears showed that PfAPR1-KO parasites progressed to the late schizont stage but, unlike WT, their released merozoites remained adjacent to the surface of neighboring RBCs rather than invading **(Fig. 2b)**. We further monitored schizont egress and invasion by collecting samples every four hours from 44 to 56 hpi using a conventional Hemacolor staining. This analysis revealed that PfAPR1-KO parasites exhibited normal egress but failed to invade new RBCs and remained adjacent to their surface, in contrast to the robust invasion observed in controls **(Fig. 2c)**. Collectively, these results demonstrate that PfAPR1 is essential for asexual parasite replication, specifically for the merozoite invasion of RBCs.

**Fig. 2:**
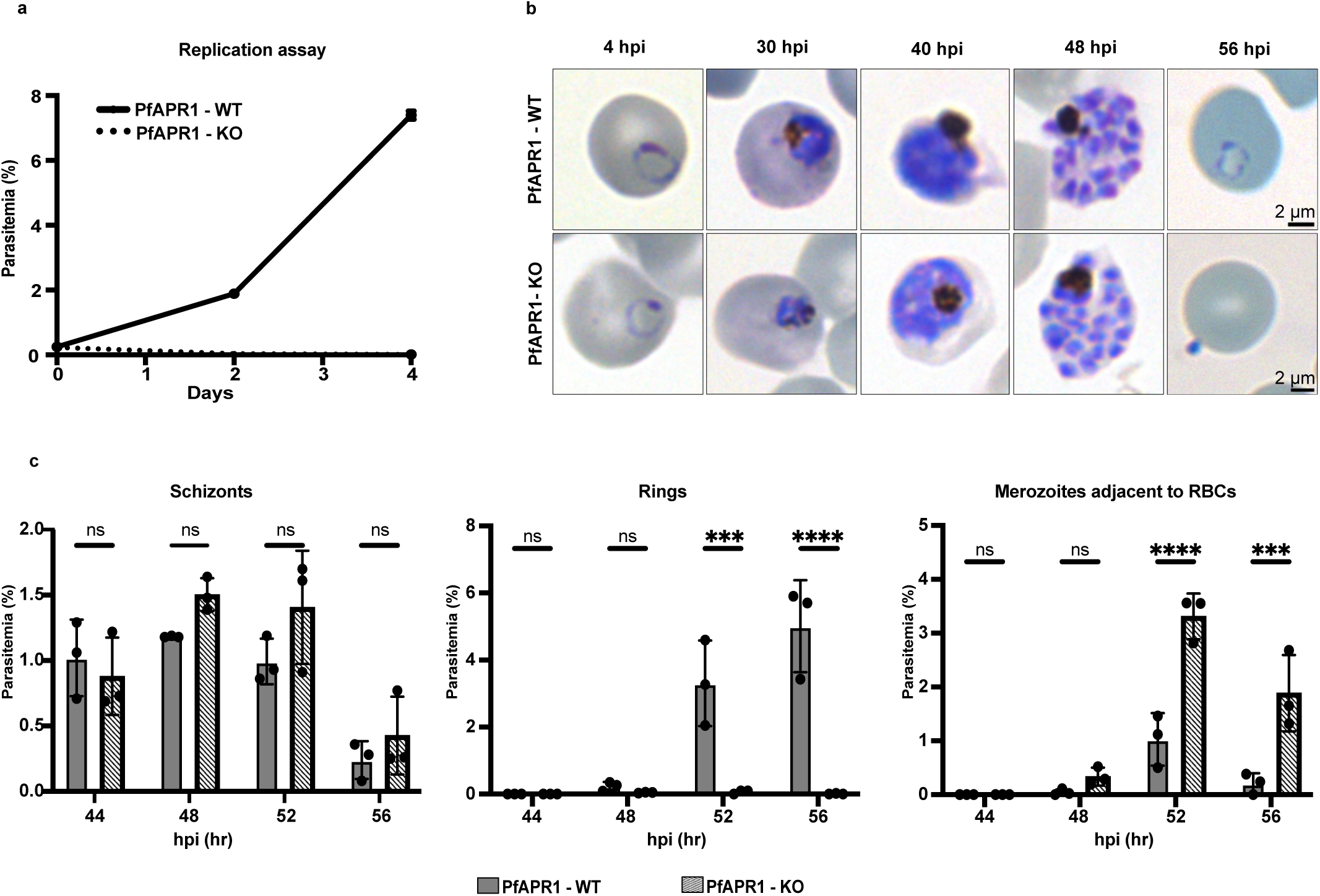
PfAPR1 is essential for asexual parasite replication. **a,** Synchronized PfAPR1^DiCre^parasites were treated with either DMSO or rapamycin, and their growth was monitored over two consecutive replication cycles by flow cytometry (n=3, error bar represents standard deviation). PfAPR1-KO parasites (+rapa) exhibited a lethal growth defect. **b,** Representative Hemacolor-stained smears of PfAPR1^DiCre^parasites were prepared for during the first replication cycle of rapamycin or DMSO treatment. PfAPR1-KO parasites progressed similarly to PfAPR1-WT parasites until the late schizont stage but were unable to invade new RBCs and remained attached to the RBC surface. Scale bars: 2 µm. **c,** Tightly synchronized PfAPR1^DiCre^ parasite were smeared and stained with Hemacolor every four hours from 44 to 56 hpi and parasitemia was quantified by light microscopy. PfAPR1-KO (+ rapa) parasites showed no significant difference in schizont development or egress; however, mature merozoites failed to form new rings and remained attached to RBCs. Data represent three independent biological replicates, error bar represents ±standard deviation (SD). Statistical significance was determined by two-way ANOVA: ****p <0.0001; ***p <0.0002; ns, not significant (p ≥ 0.3179).

### PfAPR1 is dispensable for IMC formation but modulates SPMT organization

APR proteins organize the cytoskeletal structure - including the IMC and SPMTs - and maintain apical polarity in *T. gondii* and *Plasmodium spp*.^4,16,26,28,39,40^. To determine whether PfAPR1 contributes to cytoskeletal organization in *P. falciparum*, we analyzed PfAPR1-KO parasites using U-ExM. Loss of PfAPR1 did not disrupt IMC formation. However, all PfAPR1-KO schizont examined displayed at least one multinucleated, morphologically abnormal daughter cell enclosed by IMC, a phenotype not observed in WT parasites **(Fig. 3a)**. Thus, PfAPR1 is not required for IMC formation, but it likely contributes, either directly or indirectly, to the fidelity of daughter cell segmentation.

**Fig. 3:**
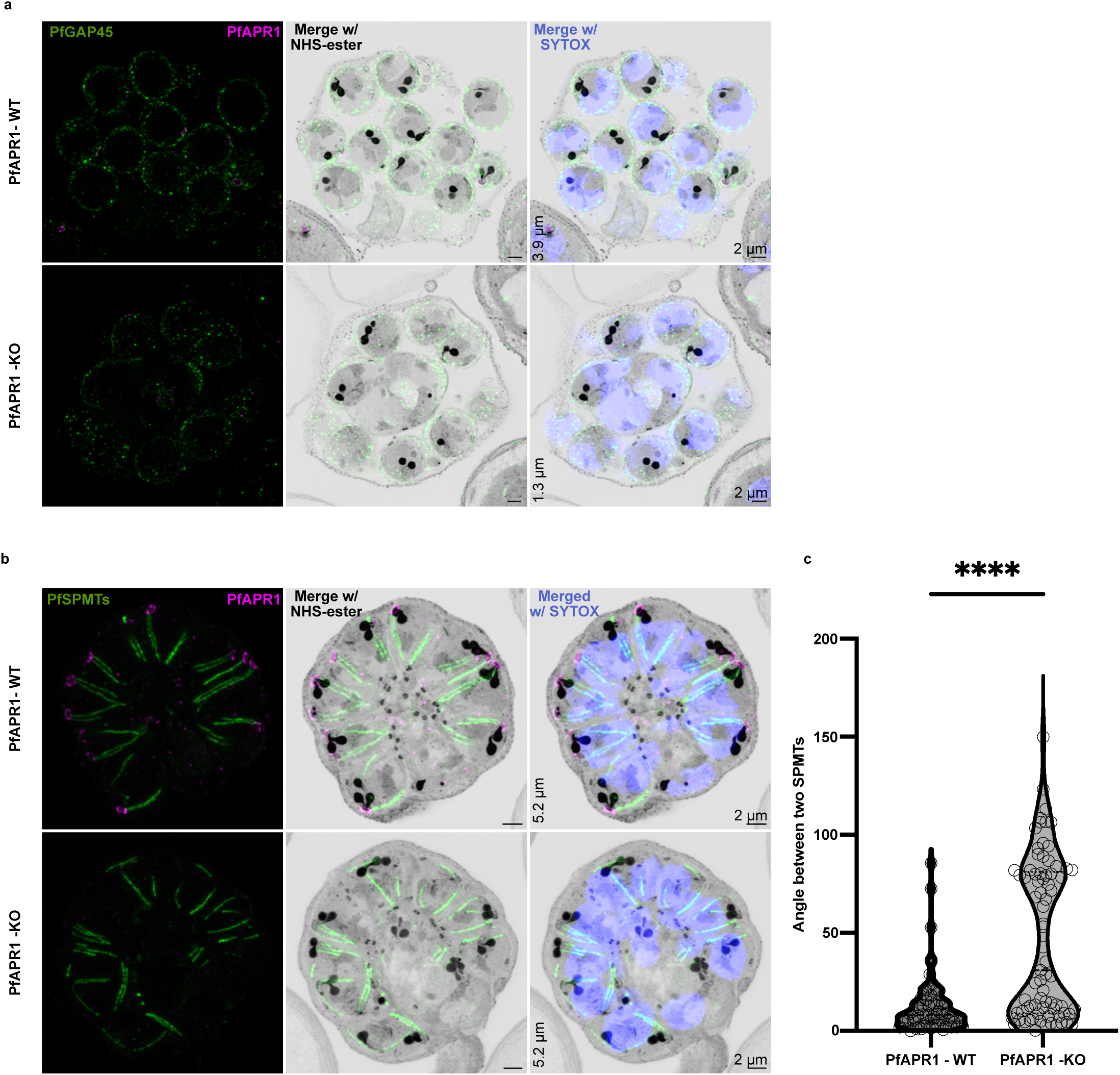
PfAPR1 is dispensable for IMC formation and maintaining apical polarity but required for daughter cell segmentation and SPMT organization. **a,** U-ExM images of E64-arrested PfAPR1^DiCre^ parasites probed with ɑ-PfGAP45 (green) show that loss of PfAPR1 does not affect IMC formation, as daughter cells remain surrounded by IMC in both WT and KO parasites. However, multinucleated, morphologically abnormal daughter cells are consistently observed in PfAPR1-KO (+rapa) parasites, compared to controls. **b and c,** ML10-stalled PfAPR1^DiCre^ parasites were probed for SPMTs (ɑ-tubulin, green). U-ExM reveals that, while SPMTs form and remain associated with the NHS-ester-stained APR in both WT and KO parasites, more than half of daughter cells in PfAPR1-KO parasites display disparate or non-parallel SPMTs. The angle between two SPMTs in control and PfAPR1-KO parasites was calculated for each SPMT using the Neuroanatomy plugin in FIJI. The violin plot represents data from one biological replicate, with individual data points shown as circles within the plot; additional replicates are shown in the **Extended Data Fig. S3a**. For each condition, three schizonts per replicate and at least 20 SPMT pairs per schizont were analyzed. Statistical analysis was performed using Welch’s two tailed unpaired t-test ( p < 0.0001). All U-ExM images are representative of a minimum of three independent biological replicates. NHS-ester (grey) is used as a general protein stain, and SYTOX is used to stain nuclei in U-ExM. Scale bars: 2 µm. For U-ExM images, z-stack depth is indicated in µm.

To further assess SPMT formation and organization, mature parasites were arrested prior to parasitophorous vacuolar membrane (PVM) rupture with the PKG-inhibitor, ML10^41^, and stained with an ɑ-tubulin antibody. In both WT and PfAPR1-KO parasites, SPMTs formed and remained associated with the NHS-ester-stained APR. Consistent with previous reports^42,43^, WT parasites displayed paired, nearly parallel SPMTs aligned along one side of each merozoite, with an average angle of 14.3° ± 1.48° between the two SPMTs. In contrast, PfAPR1-KO schizonts exhibited a significantly increased average angle of 47.3° ± 1.73° between SPMTs, indicating a loss of this parallel alignment. Although some PfAPR1-KO daughter cells preserved the parallel SPMT arrangement, more than 50% displayed disorganized or non-parallel alignment **(Fig. 3b and c, Extended Data Fig. S3a )**.

Furthermore, NHS-ester-staining confirmed that apical polarity was maintained in PfAPR1-KO parasites, with rhoptries apically positioned, and the basal complex ring at the basal end of developing merozoites **(Fig. 3a and b)**. Importantly, the dense APR structure was preserved, indicating that loss of PfAPR1 alone is not sufficient to disrupt overall cytoskeletal architecture. Together, these results indicate that while PfAPR1 is dispensable for apical polarity and IMC formation, but it is important for the fidelity of daughter cell packaging and SPMT organization during schizogony.

### PfAPR1 severely affects invasion, particularly after pre-invasion step

To define the role of PfAPR1 during invasion, we analyzed synchronized PfAPR^DiCre^ parasites (± rapamycin) immediately after release from ML10-induced arrest, using IFA, live-cell differential interference contrast (DIC) microscopy, and U-ExM. Merozoite invasion of RBCs is a rapid, multi-step process lasting 5-10 min of contact and can be divided into pre-invasion (attachment and reorientation), internalization (tight junction formation and entry) and ring-stage establishment^44–46^. During invasion, merozoites target the RBC cytoskeleton to modulate membrane deformability; pre-invasion causes mild deformation, while internalization induces pronounced, reversible echinocytosis, characterized by a spiky membrane morphology^46,47^. Time-lapse DIC microscopy revealed that WT merozoites efficiently attached, reoriented, invaded, and established ring-stage infection, accompanied by characteristic RBC deformations and reversible echinocytosis **(Fig. 4a, Supplementary Video S1)**. In contrast, PfAPR1-KO parasites attached and reoriented on the RBC but failed to invade, often bouncing on the RBC surface before falling off. These mutants induced mild deformation and membrane wrapping, with RBCs displaying abnormal, thin needle-like protrusions rather than typical echinocytosis **(Fig. 4a, Supplementary Video S2)**. Quantification of invasion efficiency per RBC contact of WT parasites showed that, 138 of 167 (82.63%) WT merozoites contacting RBCs, successfully invaded. In stark contrast, only 3 of 68 (4.4%) PfAPR1-KO merozoites invaded, with the remaining 65 (95.6%) failed to do so **(Fig. 4b)**. This severe invasion defect closely resembles the phenotype observed with R1 peptide inhibition of the PfAMA1-RON complex^48–50^, suggesting that PfAPR1 is critical for invasion steps following merozoite reorientation.

**Fig. 4:**
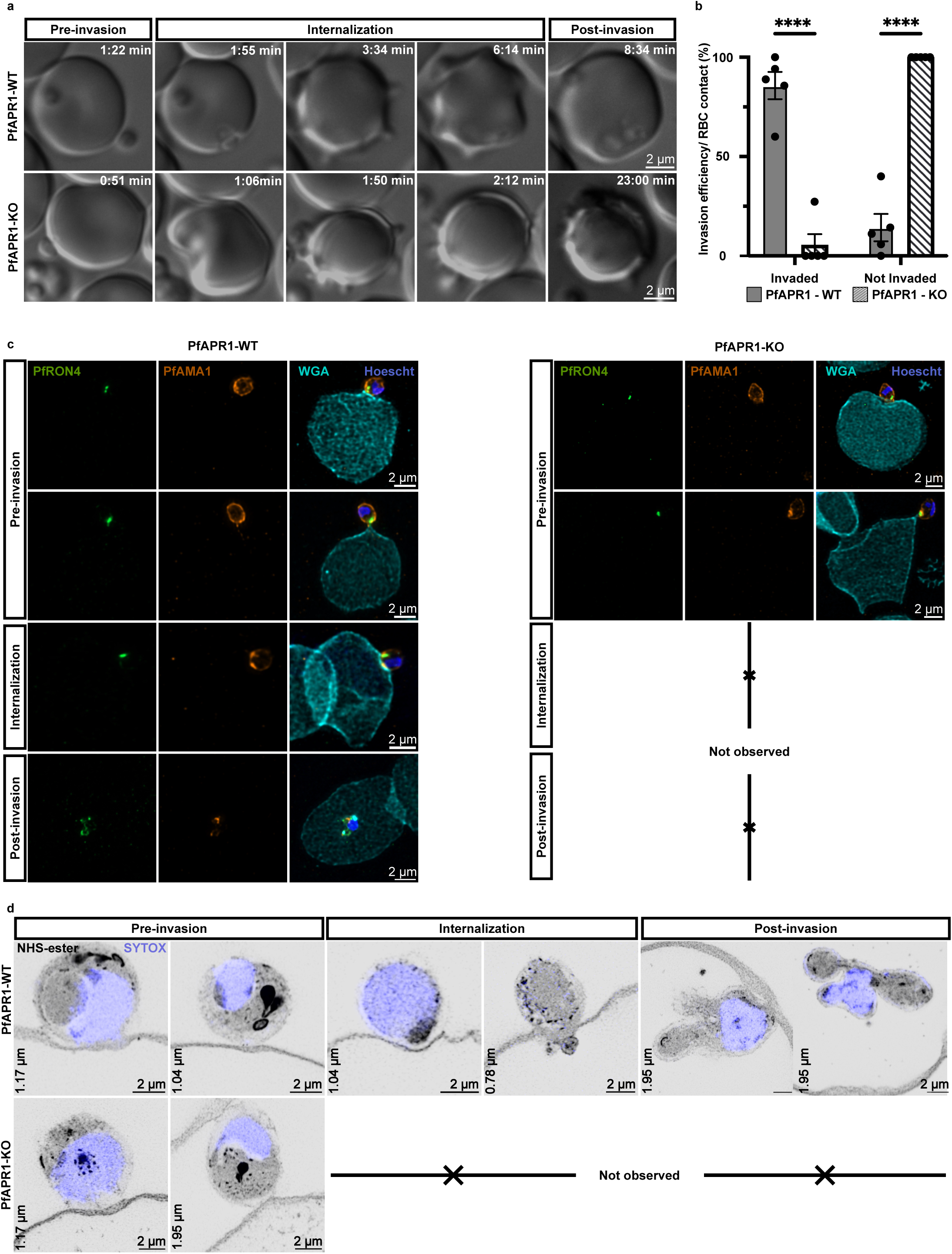
PfAPR1 is required for tight junction formation and invasion in *P. falciparum.* Synchronized PfAPR1^DiCre^ parasites (± rapamycin) were Percoll-purified and incubated with 25 nM ML10 for 2 h prior to egress. After washing, schizonts were mixed with fresh RBCs to perform invasion assays. **a and b,** Differential interference contrast (DIC) live-cell imaging showed that PfAPR1-KO parasites could contact RBCs but failed to invade, whereas control parasites successfully invaded RBCs. Data represent five independent biological replicates. Error bars indicate mean ± s.e.m. Statistical analysis was performed using two-way ANOVA. **c,** IFA images (maximum projection) demonstrate that PfAMA1 (orange) and PfRON4 (green) form a tight junction in WT parasites, but not in PfAPR1-KO parasites during invasion. **d,** High-resolution NHS-ester-stained U-ExM images show that WT parasites progress through all stages of invasion, while PfAPR1-KO parasites were only observed at pre-invasion stages, including attachment and reorientation of the apical tip of merozoite. Nuclei were labeled with Hoechst in IFA and with SYTOX in U-ExM. NHS-ester staining was used to visualize general protein content. Images shown are representation of minimum of five independent replicates.

Given this similarity, we next investigated whether PfAPR1 ablation affects PfAMA1 expression or trafficking. PfAMA1 is a microneme protein, which is proteolytically processed and translocated to the merozoite surface during egress, a prerequisite for its role in invasion^37,51^. IFA with ɑ-PfAMA1 antibodies on mid- and E64-arrested schizonts revealed normal microneme expression and surface translocation in both WT and PfAPR1-KO parasites **(Extended Data Fig. S3b)**. This indicates PfAPR1 does not affect PfAMA1 expression or spatial redistribution (i.e., translocation). Thus, the invasion defect likely occurs downstream of PfAMA1 surface exposure, potentially in post-reorientation steps involving the PfAMA1/PfRON complex. To investigate this, we tracked PfRON4 and PfAMA1 localization during invasion. PfRON4 is a rhoptry neck protein and component of the PfRON complex, interacts with PfAMA1 to form tight junction^45,52,53^. In WT parasites, PfRON4 redistributes from the apical rhoptry neck to an arc or ring-like configuration, observed as paired puncta in single z-slices, corresponding to the developing tight junction and then disperses along the PVM following invasion. In PfAPR1-KO parasites, PfRON4 remains confined to a single apical focus throughout merozoite attachment and reorientation, with no evidence of the characteristic redistribution or tight junction formation **(Fig. 4c)**. These results suggests that PfAPR1 is required for tight junction formation during invasion.

To visualize ultrastructural defects during invasion, we employed U-ExM with NHS-ester staining. In WT parasites, U-ExM captured all invasion stages, including internalization. In contrast, PfAPR1-KO parasites exhibited persistent attachment and reorientation, without evidence of internalization **(Fig. 4d, Extended Data Fig. S3c and d)**. Collectively, these results demonstrate that PfAPR1 is essential for invasion steps following merozoite reorientation, likely by enabling the redistribution of rhoptry neck proteins such as PfRON4 - a prerequisite for tight junction formation and successful invasion. Our findings confirm PfAPR1 as a key regulator of post-reorientation events during RBC invasion.

### PfAPR1-interacting proteins (PfAPR4, PfCHAKRA, and PfBBx)

The persistence of NHS-ester-stained APR structures in PfAPR1-KO parasites suggested the presence of additional structural proteins. To identify PfAPR1 interactors, we conducted a co-immunoprecipitation (co-IP) in PfAPR1^DiCre^ schizonts (40 - 48hpi) using an α-HA antibody and parental PfNF54^DiCre^ parasites as a control. Unbiased data-dependent acquisition (DDA) mass spectrometry identified 49 high confidence interactors, (≥ 10-fold enrichment in experimental samples and ≥ 10 unique peptides across replicates (**Supplementary Table 1**). We prioritized three uncharacterized proteins - PF3D7_1128900, PF3D7_1138000, and PF3D7_1307900 **(Fig. 5a)**. These proteins are predicted to be essential for parasite replication based on transposon screens^30,31^. PF3D7_1128900 (73.8 kDa) is a conserved protein with unknown function and no predicted domains. The *T. gondii* ortholog (TgAPR4) of PF3D7_1128900 localizes to the APR^4,26^, thus we named this protein **PfAPR4**. PF3D7_1128900 is a 335.7 kDa conserved *Plasmodium*-specific protein with unknown function. We propose to name it as **C**ytoskeletal **H**ook **A**pi**c**al **K**ey **R**ing-**A**ssociated protein (**PfCHAKRA**). PF3D7_1307900 (105 kDa) is a putative tripartite motif protein, which contains a B-box-type zinc finger domain (IPR000315)^54^ and is designated as **B**-**b**o**x** protein, thus we have named it **PfBBx**. To determine their localization, we generated a dual-transgenic parasite strain (PfGOI^V5/HA^) by fusing a spaghetti monster V5 (smV5) tag to the 3’ end of each gene in the PfAPR1^DiCre^ background (or the NF54^DiCre(*pfs47*)^ for PfBBx fused with smHA tag) **(Extended Data Fig. S4a-d)**. Since PfAPR1 is primarily expressed in schizonts, we performed IFA with ɑ-V5 antibodies at this stage. IFA demonstrated that all three candidates localized as ring-like structures at the apical apex of the IMC marker, PfGAP45 **(Fig. 5b)**. Super-resolution U-ExM further confirmed their localization aligned with NHS-ester-stained APR, consistent with PfAPR1’s localization **(Fig. 5c)**.

**Fig. 5:**
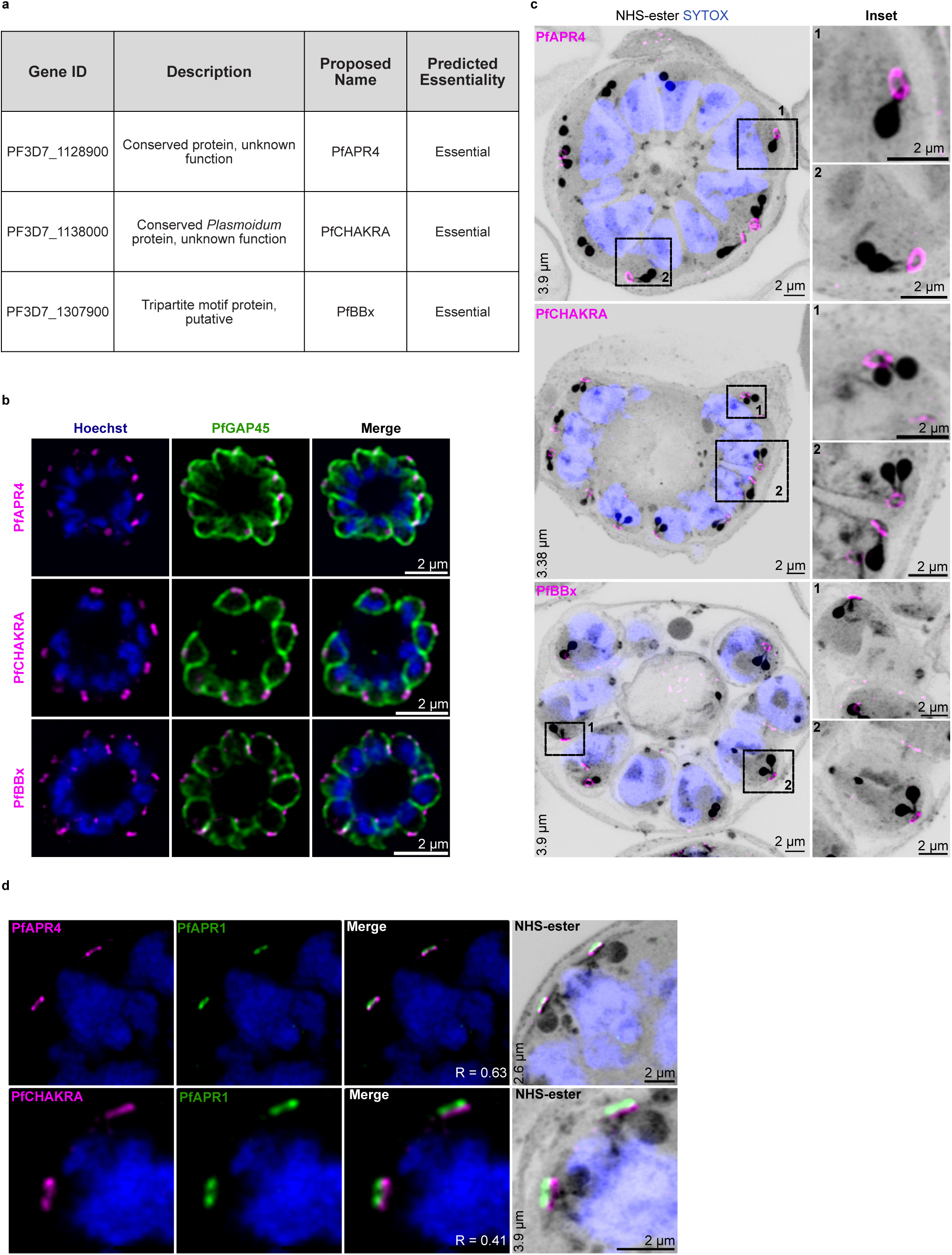
Identification and localization of PfAPR1-interacting partners. Schizont-stage PfAPR1^DiCre^ parasites (experimental sample) and PfNF54^DiCre^ (control) were subjected to co-immunoprecipitation (co-IP) by ɑ-HA antibody (two independent biological replicates). Supplementary Table 1 lists candidate PfAPR1-interacting proteins selected based on two criteria: (1) ≥ 10-fold enrichment in experimental samples and (2) ≥ 10 unique peptides across replicates. **a,** Gene ID and annotations for three selected candidates (PfAPR4, PfCHAKRA, and PfBBx), including predicted essentiality from transposon mutagenesis screen^31^ and new given name. Transgenic parasite strains expressing identified proteins in PfAPR1^DiCre^ background (or PfNF54^DiCre^ for PfBBx) were generated and harvested at schizont stage for IFA and U-ExM. **b,** IFA images (single z-stack slice) showing a ring-like structures formed by PfAPR4, PfCHAKRA, and PfBBx (ɑ-V5 or HA, magenta) at the apical end of PfGAP45 (IMC marker, green) in schizonts. **c,** High-resolution U-ExM images reveal that ring-like pattern of all three proteins colocalize with NHS-ester-stained APR. **d,** U-ExM demonstrates PfAPR4 and PfCHAKRA (ɑ-V5, magenta) localize beneath PfAPR1 (ɑ-HA, green), with PfCHAKRA positioned proximal to the proto-APR structures, stained by NHS-ester. Pearson’s R values (calculated by Coloc2 plugin in Fiji) are shown in the right corner. Nuclei were stained with Hoechst (IFA) or SYTOX (U-ExM). NHS-ester provided general protein staining in U-ExM. Images are representative of three independent biological replicates. Boxed regions are magnified in insets. Scale bar, 2 µm. z-depth for U-ExM is indicated (left corners).

To assess the colocalization with PfAPR1, we analyzed PfAPR4 and PfCHAKRA in the dual-transgenic parasites using U-ExM. U-ExM revealed that PfAPR4 and PfCHAKRA (ɑ-V5) are localized beneath PfAPR1 (ɑ-HA), with PfAPR4 in closer proximity (Pearson’s R = 0.63) than PfCHAKRA (R = 0.41) **(Fig. 5d, Extended Data Fig. S4e)**. These results confirm that three PfAPR1-interacting proteins are all additional APR residents, though their functional roles remain to be characterized.

### APR biogenesis in *P. falciparum*

While APR assembly in *T. gondii* proceeds from centrosome-associated arcs to a mature ring structure^27^, the corresponding process in *Plasmodium* remains poorly understood. Using U-ExM with NHS-ester staining, we tracked the localization of PfAPR1 and PfCHAKRA during schizogony. The NHS-ester efficiently labels the protein-dense APR, its precursor (proto-APR), and the centriolar plaque (CP)^35^. The CP is a non-conventional centrosome, which forms a bipartite structure with distinct extranuclear and intranuclear regions. The extranuclear region is hypothesized to nucleate cytoplasmic sub-pellicular microtubules and is physically connected to the proto-APR^35,55^, while the intranuclear region is a chromatin-free area that serves as the nucleation site for nuclear microtubules^56–58^. Notably, the CP duplicates once per nuclear cycle, becoming less visible at the end of the schizogony^35,59–61^. Our U-ExM revealed that both PfAPR1 and PfCHAKRA are recruited to the apical end of the extranuclear CP, forming an arc-shaped structure that coincide with the NHS-ester stained proto-APR. Notably, PfCHAKRA is positioned closer to the proto-APR than PfAPR1, suggesting spatial organization during APR assembly. As the CP duplicated during the nuclear cycle, we observed that proto-APR along with PfAPR1, and PfCHAKRA also duplicate, forming paired semi-circles. Notably, the maturation into the closed APR rings occurred asynchronously; one duplicated CP exhibited a closed ring-shaped APR, while the other retained an arc-shaped proto-APR. These structures subsequently mature into the closed ring-shaped APRs in the newly formed CPs. At the end of schizogony, when the CP disappears, the NHS-ester-labelled APR ring persisted at the apical end of mature daughter cells, with PfAPR1 remaining associated, whereas PfCHAKRA was no longer visible **(Fig. 6, Extended Data Fig. S5)**. Collectively, these results demonstrate that APR biogenesis in *Plasmodium* involves a conserved arc-to-ring maturation, with PfAPR1 and PfCHAKRA occupying distinct spatial positions and highlights the close relationship between CP duplication and APR assembly.

**Fig. 6:**
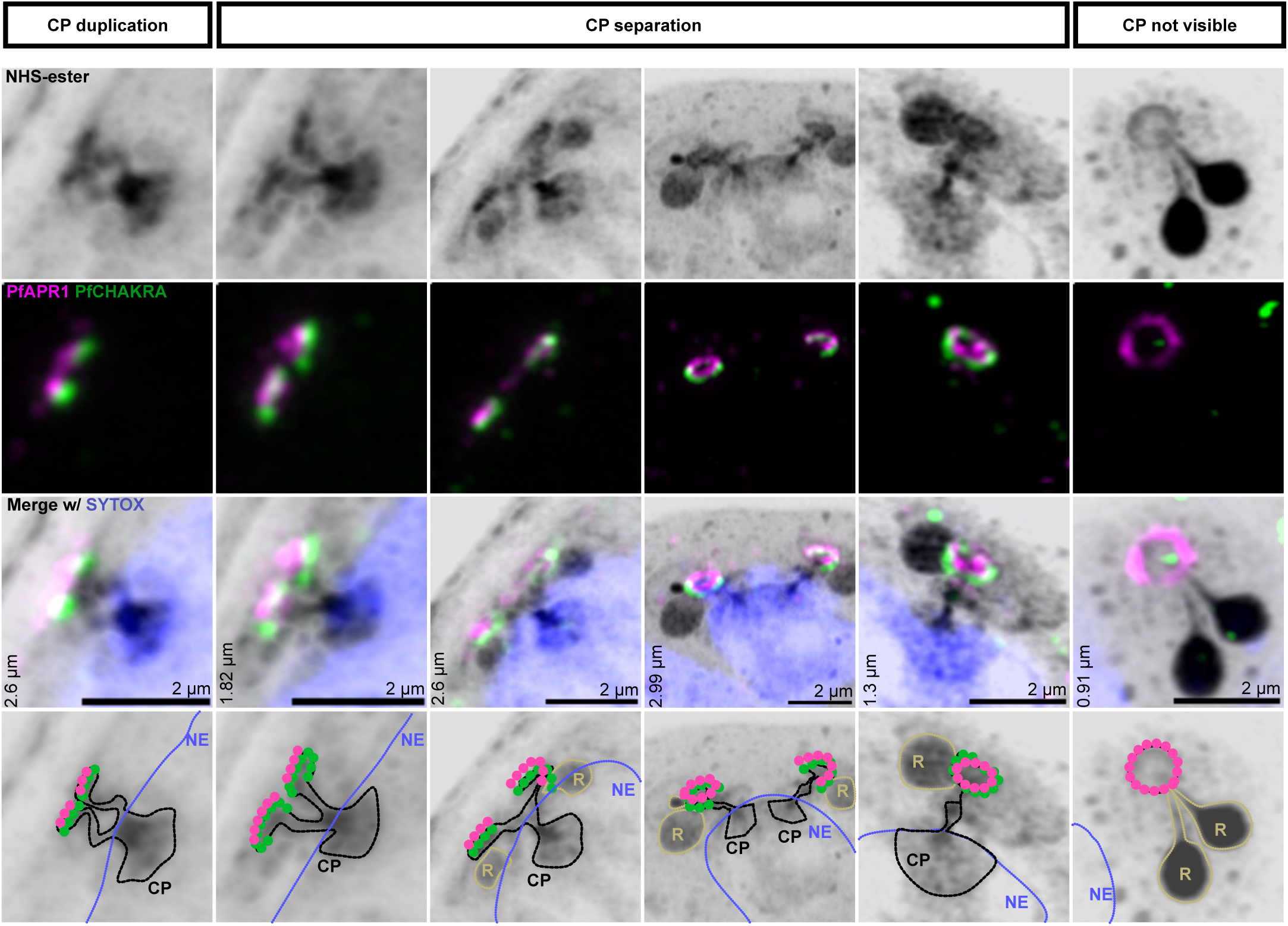
APR biogenesis in *P. falciparum* visualized by U-ExM in dual transgenic PfCHAKRA^V5^ parasites: U-ExM was used to monitor the dynamic expression and localization of PfAPR1 and PfCHAKRA during schizogony. First row: NHS-ester staining highlights the densely stained centriolar plaque (CP) and APR. Second row: PfAPR1 (ɑ-HA, magenta) and PfCHAKRA (ɑ-V5, green) localizes to the extranuclear CP region, coinciding with nascent and mature APR structures. APR formation progresses from an arc to a closed circle. Third row: Merged images of panel 1 and 2 provide combined visualization of protein localization and sub-cellular structures. Fourth row: Schematic diagram illustrating APR biogenesis, interpreted from U-ExM data. Images are representative of three independent biological replicates. Z-depth is indicated on the left corner of each image. Scale bar = 2 µm.

## Discussion

The APR is a central feature of the apical complex in all apicomplexans and plays a critical role in parasite shape, daughter cell segregation during cell division, and host cell invasion^6,10,23^. However, the molecular composition and functional role of the APR in *P. falciparum* has remained elusive. To address this gap, we identified and functionally characterized PfAPR1 as the first APR-resident protein in the asexual stage of *P. falciparum*. PfAPR1 is essential for asexual replication, with a complete block of merozoite invasion following inducible deletion. Using PfAPR1 as a molecular bait, we identified three additional APR-resident proteins in *P. falciparum*: PfAPR4, PfCHAKRA, and PfBBx.

In this study, we leveraged advanced microscopy approaches to resolve the intricate architecture, assembly, and function of the APR in *P. falciparum*. U-ExM has emerged as a transformative tool in apicomplexan research, enabling visualization of subcellular structures at a resolution previously attainable only by electron microscopy^10,35^. Importantly, we extended the application of U-ExM beyond static organization of the APR by integrating it with dynamic invasion assays, allowing us to capture the sequential stages of merozoite invasion - attachment, reorientation, and entry into RBCs - at high resolution. NHS-ester staining further highlighted the protein-dense APR regions and enabled tracking of dynamic changes during the invasion stages. Building on these advances, we adapted and implemented iterative U-ExM (iU-ExM)^62–64^, achieving ∼15-fold linear expansion, to overcome the resolution limits of standard U-ExM. While standard U-ExM allowed us to readily resolve the APR as a single ring in merozoites, iU-ExM enabled us to distinguish two separate rings and accurately localize PfAPR1, a level of detail previously only possible with electron microscopy or electron tomography^6,9,43,65–67^. We speculate that the two apical rings are likely less protein-dense compared to the basal ring, potentially explaining their limited visibility with both standard and iU-ExM using NHS-ester staining. This result is supported by recent cryo-ET analysis, which reveal that the basal ring is 50 nm thick and amorphous, while the upper two rings display distinct periodicity but sometimes variable geometry appearing bent or distorted^6^. Our approach enables these insights using standard fluorescence microscopy, obviating the need for highly specialized EM facilities. To our knowledge, this is the first application of U-ExM to study the invasion process and of iU-ExM to dissect a complex structure within *P. falciparum*.

Our findings demonstrate that the loss of PfAPR1 does not catastrophically disrupt the formation of the cytoskeletal network (IMC and SPMTs) in *P. falciparum*, but it does cause mild defects in cell segmentation and SPMT organization. Previous studies have hypothesized the APR to be an MTOC for SPMTs. However, our data and recent studies in *Plasmodium spp.* and *T. gondii* suggest that the APR mainly anchors SPMTs rather than nucleating them^16,26,27,39^. In PfAPR1-KO merozoites, the apical ends of SPMTs remain closely associated with the NHS-ester-dense APR, but notably, more than 50% of merozoites display SPMT disorganization or non-parallel SPMT arrangement, characterized by loss of typical alignment between SPMTs. This pronounced phenotype suggests that proper SPMT organization is highly dependent on PfAPR1, even in merozoites, which possess only 2 - 4 SPMTs. While our analysis is limited to the merozoite stage, the potential role of PfAPR1 in other zoite stages with a higher number of SPMTs and more complex cytoskeletal network such as the ookinete remains to be explored.

Furthermore, our data show that the APR structure persists in PfAPR1-KO parasites, indicating that while PfAPR1 is a structural component, other proteins also contribute to APR integrity. This is consistent with findings in *T. gondii*, where overlapping functions among APR-associated proteins help maintain structure; for example, loss of both TgAPR1 and TgKinesinA disrupts the APR, but depletion of TgAPR1 alone has only modest effects^39^. Collectively, these findings highlight the cooperative roles of APR components in maintaining cytoskeletal architecture and enabling host cell invasion, with PfAPR1 contributing to, but not solely defining, APR stability.

Through live-cell microscopy, we observed that PfAPR1-KO parasites egress from RBCs normally and complete pre-invasion steps, including attachment and apical reorientation, but repeatedly fail to invade new RBCs and eventually fall off the RBC surface. IFA analysis with PfRON4 and PfAMA1 confirmed that, although PfAPR1-KO parasites can attach and reorient, they are unable to form a functional tight junction, a step that is essential for successful invasion. Notably, the RBCs in contact with PfAPR1-KO merozoites exhibited atypical echinocytosis, characterized by transformation into a rounded sphere with numerous needle-like protrusions. This phenotype closely resembles that induced by R1 treatment, which blocks AMA1-RON4 interaction and tight junction formation^44,68^. Both R1 peptide treatment and PfAPR1-KO parasites induce initial RBC deformation and pronounced atypical echinocytosis, suggesting that initial contact and attempted invasion are sufficient to trigger these membrane changes, even in the absence of successful entry into RBCs. Taken together, our data suggest that PfAPR1 is required for tight junction formation or stabilization, and that its absence phenocopies the effects of direct PfAMA1-RON complex inhibition. Further studies are needed to elucidate the molecular role of PfAPR1 in tight junction assembly and its potential regulation of invasion-induced RBC membrane deformation.

In summary, using reverse genetics and advanced microscopy techniques, our study establishes that PfAPR1 is a pivotal structural and functional component of the APR in *P. falciparum*. PfAPR1 is essential for maintaining cytoskeletal organization and enabling RBC invasion. By elucidating the role of PfAPR1 and identifying additional APR-resident proteins, we provide new insights into the molecular composition of this critical APR structure. These findings not only advance our understanding of APR biology but also highlight the APR as a promising target for developing next-generation antimalarial strategies.

## Methods

### Plasmid construction and generation of transfected lines

#### PfAPR1^DiCre^ (pPG35)

The PfAPR1^DiCre^ parasite strain was generated using a donor plasmid assembled from five PCR-amplified DNA fragments via Golden Gate (GG) assembly with the NEB BsaI-HFv2 kit (E1601L, NEB), following the manufacturer’s protocol. The fragments included: (1) the 5’ UTR region, amplified from 3D7 genomic DNA with primers oJDD7410/7411; (2) a codon-optimized PfAPR1 gene, amplified from the synthetic gene block GB85 (Biobasic) using oJDD7412/7413; (3) a region containing the linker-3xHA, LoxP, and human dihydrofolate reductase (hDHFR) amplified from pPG31 with oJDD7414/7415; (4) the 3’ UTR region, amplified from 3D7 gDNA with oJDD7416/7417; (5) the pGEM ampicillin region, amplified from plasmid pPG25 using oJDD7418/7419.

#### PfAPR4^V5^ (pPG61)

The PfAPR4^V5^ parasite strain was produced by transfecting the donor plasmid pPG61 into the PfAPR1^DiCre^ parasite. The pPG61 plasmid was assembled from five PCR-amplified fragments using GG assembly. The fragments comprised: (1) a 5’ homology region (HR) amplified from 3D7 genomic DNA with oJDD8357/8358, and a codon-optimized PfAPR4 gene from gene block GB100 with oJDD8463/8464 (joined by PCR sewing); (2) smV5, amplified from pPG03 with oJDD8331/8332; (3) blasticidin S deaminase (BSD) cassette, amplified from pPB69 with primers oJDD8333/8334; (4) a 3’ HR region amplified from 3D7 genomic DNA with oJDD8361/8362; and (5) the pGEM backbone, amplified from pPG25 with oJDD8337/8338.

#### PfCHAKRA^V5^ (pPG72)

The PfCHAKRA^V5^ parasite strain was generated by integrating the pPG72 construct into the PfAPR1^DiCre^ parasite background. The pPG72 plasmid was assembled from five PCR-amplified fragments using a GG reaction. Piece 1 was obtained by PCR sewing a 5’ HR region from 3D7 genomic DNA (oJDD8375/8376) with a codon-optimized region from gene block GB102 (oJDD8466/8378); piece 2 was smV5, amplified from pPG03 with oJDD8331/8332; piece 3 comprised BSD, amplified from pPB69 with primers oJDD8333/8334; piece 4 was 3’ HR from 3D7 genome with oJDD8379/8380; and piece 5 was the pGEM backbone, amplified from pPG25 with oJDD8337/8338.

#### PfBBx^HA^(pIA39)

The pIA39 was generated by assembling four PCR-amplified DNA fragments using GG assembly. The fragments included: (1) a 5’ HR region amplified from 3D7 gDNA with oJDD8875/8876; (2) smHA and BSD, amplified from pPB69 with oJDD8877/8878; (3) a 3’ HR region amplified from 3D7 genomic DNA with primers oJDD8879/8880; and (4) the pGEM backbone, amplified from pPB69 with oJDD8818/8881. The assembled construct was used for transfection to generate the PfBBx^HA^ parasite in PfNF54^DiCre^ background.

All CRISPR-Cas9 guide plasmids were generated by annealing and ligating oligonucleotides corresponding to guide sequences into the BpiI-digested pRR216 plasmid. The pRR216 plasmid contains SpCas9 and a U6 promoter-driven guide RNA cassette^69^. For PfAPR1^DiCre^ parasites, plasmid pPG36 with guide oligos oJDD7420 and oJDD7421 was used. For PfAPR4^V5^ parasites, guide plasmids pPG62 (oJDD8403/8404), pPG63 (oJDD8405/8406), and pPG64 (oJDD8407/8408) were used. For PfCHAKRA^V5^ parasites, guide plasmids - pPG73 (oJDD8419/8420) and pPG74 (oJDD8421/8422) were used. For PfBBx^HA^ parasites, pIA44 (oJDD8578/8579) and pIA45 (oJDD8580/8581) were used as guide plasmids.

### Parasite culture and maintenance

The *P. falciparum* NF54^DiCre(pfs47)^ laboratory strain, obtained from Moritz Treeck^34^, was used throughout this study as the parental background for transfections and as a control strain. Parasites were cultured in RPMI-1640 medium (Sigma, R6504) supplemented with 25 mM HEPES [4-(2-hydroxyethyl)-1-piperazineethanesulfonic acid] (EMD Biosciences), 0.21% sodium bicarbonate (VWR, 0511), 50 mg/L hypoxanthine (Sigma), and 0.5% Albumax II (Invitrogen). Human O^+^ erythrocytes, obtained from Valley Biomedical, were used for parasite culture. Cultures were maintained at 37°C under a gas mixture of 5% CO_2_, 1% O_2_, and 95% N_2_. Parasite cultures were synchronized using the Percoll-sorbitol method, as previously described^70,71^. Briefly, tightly synchronized ring-stage parasites were obtained by subjecting late-stage schizonts to a 60% Percoll (GE Healthcare), followed by sorbitol lysis at 0-4 hpi. This process was repeated as needed to ensure synchronization within a 2–4 h window.

### Transgenic parasite generation

To generate transgenic parasite strains (PfAPR1^DiCre^, PfAPR4^V5^, PfCHAKRA^V5^, and PfBBx^HA^), 25 μg of linearized and purified donor plasmid DNA and 25 μg of SpCas9-plasmid DNA containing the appropriate gRNA sequence were co-transfected into Percoll-purified late schizonts (either NF54^DiCre(pfs47)^ or PfAPR1^DiCre^ background) using a P3 Primary cell 4D-Nucleofector X Kit L (Lonza, V4XP-3024) and Amaxa 4D electroporator (Lonza, program FP158). After electroporation, parasites were immediately transferred to fresh RBCs at 4% HCT in complete RPMI medium and returned to culture conditions described above. Drug selection was initiated 5 h post-transfection. For PfAPR1^DiCre^ parasites, 2.5 nM WR99210 (Jacobus Pharmaceuticals) was used. For PfAPR4^V5^, PfCHAKRA^V5^, and PfBBx^HA^ parasites, selection was performed with 2.5 nM WR99210 (Jacobus Pharmaceuticals) and 2.5 μg/mL blasticidin (Research Products International). Parasite clones were obtained by limiting dilution in 96-well plates, and positive wells were expanded. Clonal lines were verified by PCR amplifications using gene-specific primers (listed in **Supplementary Table 2**) and by whole-genome sequencing for PfAPR1^DiCre^. For rapamycin-induced excision of target genes, tightly synchronized ring-stage parasites (0-4 hpi) were treated with 100 nM rapamycin for up to 12 h at 37°C. After induction, parasites were washed three times with pre-warmed RPMI-1640 medium to remove excess rapamycin and returned to standard culture conditions^72^.

### Reagents and antibodies

All PCR amplification were performed using the PrimeSTAR GXL DNA Polymerase kit (Takeda, R050A), according to the manufacturer’s protocol except for the elongation step which was performed at 60°C. Oligonucleotides were synthesized from Life Technologies and gene blocks were obtained from either IDT or Biobasic. The sequence of all oligonucleotides and gene blocks used in this study are provided in the **Supplementary Table 2**. A comprehensive list of all antibodies used, including their concentrations, host species, and suppliers, is presented in **Supplementary Table 3**.

### Sequence alignments

Protein sequences for PfAPR1 (*Plasmodium falciparum*: PF3D7_1141300), PbAPR1 (*P. berghei*: PBANKA_0907700), and PyAPR1 (*P. yoelii*, Py17XNL_000900117) were retrieved from PlasmoDB^30^. Sequence identity and similarity were determined using BLASTp^73^ and EMBOSS Needle^74^. Multiple sequence alignments were generated using the default CLUSTALW (MEGA 12 software^75^) and alignments were visualized with ESPript 3.0^76^.

### Replication assay

Synchronized PfAPR1^DiCre^ parasites, treated with either DMSO or 100nM rapamycin, were diluted to a final concentration of 0.25% parasitemia and 1% hematocrit. Parasite growth was monitored over two consecutive intraerythrocytic cycles. For each condition, 100 μL of culture was collected at days 0, 2, and 4 post-treatment. Samples were washed with 0.5% (w/v) BSA in PBS, resuspended in a 1:1000 dilution of SYBR green nucleic acid stain (Invitrogen #S7563), and incubated in the dark at RT for 20 min. After washing, samples were resuspended in 1x PBS. Infected RBCs were quantified using a BD FACSCalibur flow cytometer (BD Biosciences). Data were acquired with CellQuest Pro (BD Biosciences, 100,000 events per sample) and analyzed using FlowJo X software and GraphPad Prism 9. Results represent the mean ± standard deviation (SD) of three independent biological replicates.

### Western blot Analysis

Synchronized late-stage *P. falciparum* schizonts were lysed with 0.02% (w/v) saponin in PBS containing SigmaFast protease inhibitor cocktail (Sigma-Aldrich). Parasite pellets were washed with ice-cold PBS and boiled in 1x Laemmli sample buffer (Bio-Rad) at 95°C for 5 min. Protein (1x 10^8^ parasites per lane) were separated on a 4% - 20% Tris-glycine-sodium dodecyl sulfate gel (Bio-Rad) at 120 V. Proteins were transferred overnight at 30 V (4°C) to a nitrocellulose membrane (Bio-Rad) using a wet transfer system. Membranes were blocked with LI-COR Odyssey blocking buffer for 1 h at RT, then incubated with primary antibodies (α-HA, 1:1000; α-LDH, 1:1000) and IRDye-conjugated secondary antibodies. Blots were imaged on a LI-COR Odyssey CLx imaging system, and fluorescence signal intensities were quantified using LI-COR Image Studio 4.0 (volume measurement tool). Uncropped blot images are provided in the Source Data file.

### Immunofluorescence assays (IFA)

#### Slide-based IFA

For slide-based IFA, thin blood smears were prepared from parasite cultures and allowed to air dry completely. Smears were fixed with 4% (v/v) paraformaldehyde (PFA) in PBS for 10 min at RT, followed by three rapid washes with 1x PBS. Fixed smears were permeabilized with 0.1% Triton X-100 in PBS for 10 min at RT, then washed three times with 1x PBS for 3 min each. Non-specific binding was blocked by incubating slides in 3% (w/v) bovine serum albumin (BSA) in PBS for 1 h at RT. Slides were incubated overnight at 4°C with the respective primary antibodies diluted in 0.5% (w/v) BSA in PBS (see **Supplementary Table 3** for antibody details). The following day, slides were washed three times with 1x PBS for 5 min each. Secondary antibodies were used at 1:1000 in 0.5% BSA/PBS and, where appropriate, wheat germ agglutinin (WGA) for RBC membrane staining were applied for 45 min incubation at RT in the dark. Slides were washed three times with 1x PBS for 5 min each. DNA was stained with Hoechst 33342 (1:5,000 dilution in 1x PBS) for 20 min at RT in the dark, followed by a brief rinse with 1x PBS. Slides were prepared using Vectashield Vibrance Antifade mounting media (Vector Laboratories Inc., H-1700) and then sealed with a coverslip.

#### Batch IFA

For batch IFA, parasites were allowed to settle onto poly-D-lysine-coated coverslips placed in 24-well plates at 37°C for 20 min. The coverslips were then processed using similar fixation, permeabilization, blocking, antibody incubation, and staining steps as described for slide-based IFA.

## Image acquisition

Z-stacked images were acquired using a Zeiss LSM900 confocal microscope equipped with Airyscan 2 and a 63x oil immersion objective. Images were processed by airyscan processing or joint deconvolution using Zen software and analyzed with FIJI^77^.

## Ultrastructural Expansion Microscopy (U-ExM)

Ultrastructural expansion microscopy was performed as previously described^78^. Briefly, synchronized parasites were harvested and allowed to settle on poly-D-lysine-coated coverslips in 24-well plates for 20 min at 37°C. Parasites were fixed with pre-warmed 4 % (v/v) PFA in PBS for 20 min at 37°C, followed by three washes with pre-warmed 1x PBS. Fixed cells were then crosslinked overnight at 37°C in a solution of 1.4% (v/v) formaldehyde and 2% (w/v) acrylamide (AA) in 1x PBS. The next day, gelation was carried out by placing the coverslips in a mixture of a monomer solution (19% sodium acrylate, 10% AA, 0.1% N,N’-methylenbisacrylamide in 10x PBS), TEMED, and ammonium persulfate in a pre-chilled gelation chamber. Gelation proceeded for 5 min on ice, followed by incubation at 37°C for 1 h. After polymerization, coverslips/gels were transferred to a 6-well dish containing 1 mL denaturation buffer (200 mM SDS, 200 mM NaCl, 50mM Tris, pH 9) for 15 min with gentle agitation to detach the gels. The detached gel was then incubated with 1.5 mL denaturation buffer in a microcentrifuge tube at 95°C for 90 min to achieve complete denaturation. Gels were expanded by incubating in 25 mL deionized water (ddH_2_O) for 30 min at RT, repeated three times for optimal expansion. Gels were washed twice with 1x PBS for 15 min each at RT and blocked with 3% BSA in PBS for 30 min at RT. Small pieces of gel were cut and incubated with the respective primary antibodies diluted in 3% BSA/PBS overnight at 4°C. The next day, gels were washed thrice with 0.5% Tween-20 in PBS for 10 min each and then incubated with secondary antibodies, NHS-ester (for general protein staining), and SYTOX (for nucleic acid staining) for 2.5 h in the dark. Gels were washed again with 0.5% Tween-20 in PBS for 10 min and expanded a second time by incubation in 10 mL ddH_2_O for 30 min. Expanded gels were placed on a poly-D-lysine-coated imaging dish and imaged using a Zeiss LSM900 confocal microscope with Airyscan 2 and a 63x objective. Images were processed using Zen’s Airy-processing software and analyzed with FIJI^77^.

For colocalization analysis between PfAPR1 and its interacting partners, we manually defined a region of interest (ROI) on fluorescence images using FIJI^77^ to restrict analysis to APR region. For each ROI, colocalization between the two channels (PfAPR1 and its interacting partners) was assessed using the Coloc 2 plugin. Pearson’s correlation coefficient was calculated to quantify the degree of colocalization with Costes’ automatic thresholding used to minimize background contribution and ensure unbiased analysis.

To quantify the angle between two SPMTs in individual schizont, we used the SNT option of the Neuroanatomy plugin in FIJI for manual tracing^79^. For each selected pair of SPMTs, the first and last coordinates (x,y, and z) of each SPMT were recorded to define their direction vectors. Assuming that the SPMTs were straight lines in three-dimensional space, the angle between the two SPMTs was calculated using the following formula:

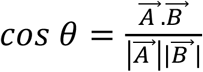

Where 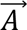 and 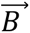 are the direction vectors of the two SPMTs. To minimize bias, file names were randomized so that SPMT tracing and measurements were performed in a manner blinded to experimental condition. Only pairs of SPMTs that were clearly visible and fully traceable throughout their length in each daughter cell were include in the analysis. The analysis was conducted across three independent biological replicates, with three schizonts examined per replicate. In each schizont, a minimum of 20 SPMT pairs were measured. All angle calculations and data analysis were performed using Microsoft Excel and GraphPad Prism.

### Iterative Ultrastructural Expansion Microscopy (iU-ExM)

Iterative ultrastructure expansion microscopy was performed as previously described^62,64,80^. PfAPR1^DiCre^ parasites were prepared as described for standard ultrastructure expansion microscopy in this study. Briefly, schizonts were purified by density gradient using 60% Percoll, settled onto glass coverslips that were coated with poly-D-lysine, and fixed with 4% paraformaldehyde. The first anchoring step was performed overnight. The first monomer solution (containing DHEBA) was added to fill the gelation chamber, followed by incubation for 15 min on ice and then 45 min incubation at 37°C. Gels were transferred to denaturation buffer and incubated at 85°C for 90 min. Gels were then expanded in water overnight (∼5-fold expansion). After expansion, gels were washed in PBS, blocked in 3% BSA, stained with primary and secondary antibodies, and re-expanded with water. Gels were then cut into approximately 1.5 cm^2^ pieces, washed three times with activated neutral gel, and incubated at 37°C for 1 h. Gels were subjected to a second round of anchoring solution overnight at 37°C. The following day, gels were washed in the third monomer solution (containing BIS) and incubated at 37°C for 1 h. Gels were then washed with 200 mM NaOH for 1 h at RT, followed by PBS wash until the pH reached 7. Gels were placed in water for a final round of expansion overnight achieving approximate ∼15-fold expansion factor. For imaging, the gel was placed on a poly-D-lysine coated glass dish and super glue was applied to the sides of the gel to prevent drift. Zeiss LSM900 confocal microscope with Airyscan 2 and a 63x objective was used to acquire images. Images were processed using Zen’s Airy-processing software and analyzed with FIJI^77^.

### Differential Interference Contrast (DIC) Live cell microscopy

For DIC live cell imaging, 0 - 2 h synchronized PfAPR1^DiCre^ parasites at 5% parasitemia and 1% hematocrit (HCT) were treated with or without 100 nM rapamycin. Parasites culture media was changed daily and Percoll-purified schizonts were arrested using 25 nM ML10 (BEI resources) for 2 h prior to egress. After arrest, parasites were washed twice with pre-warmed complete RPMI media to remove ML10. Schizonts were then mixed with fresh RBCs and allowed to settle onto a 35 mm glass-bottom imaging dish (CellView, 627871). Imaging was performed using a Zeiss LSM 980 microscope equipped with a DIC prism and a 63x oil immersion objective. The microscope chamber and imaging dish insert were pre-warmed to 37°C prior to transferring the schizonts for imaging. Parasite were imaged every 2.5 s for 30 min using DIC. A minimum of three independent biological replicates were performed.

### Affinity-based immunoprecipitation and mass spectrometry

Synchronized PfAPR1^DiCre^ and PfNF54^DiCre^ schizont cultures (300 mL each, at 3% parasitemia and 2% HCT) were harvested at 40 - 48 hpi. Parasites were released from RBC by treatment with 0.05% saponin in 1x PBS supplemented with a protease inhibitor (SigmaFast Protease Inhibitor Cocktail, Sigma). The parasite pellets obtained were resuspended in 1mL of RIPA lysis buffer (50 mM Tris-HCl, pH 7.5, 150 mM NaCl, 1% NP-40, 0.5% sodium deoxycholate, and 0.1% SDS) supplemented with protease inhibitors, and incubated for 1h at RT on a rotator. Lysates were then sonicated using a microtip sonicator at 20% amplitude for two cycles of 30s pulses, with 1 min intervals between cycles, while kept on ice. To remove hemozoin and insoluble debris, lysates were centrifuged at maximum speed (Eppendorf 5428) for 30 min at 4°C. The cleared supernatant was incubated overnight at 4°C on a rotator with magnetic beads coated with α-HA (Pierce 88836). After incubation, beads were washed three times with RIPA buffer supplemented with protease inhibitors and then washed twice with 1x PBS containing protease inhibitors. The beads were subsequently resuspended in 50 μL of 50 mM ammonium bicarbonate and sent to the Taplin Mass Spectrometry core facility at Harvard for LC-MS/MS analysis. Samples were analyzed on Thermo Orbitrap mass spectrometer using the data-dependent acquisition (DDA) mode. Summed intensities for each protein obtained in both sample and control parasites were compared, and proteins with ≥ 10 unique peptides identified across two biological replicates were considered for further analysis.

### Statistics and reproducibility

For all imaging experiments, including slide-based IFA, batch IFA, U-ExM, and live cell imaging, as well as western blot analyses, a minimum of three independent biological replicates was performed to ensure reproducibility. Immunoprecipitation experiments were conducted with two biological replicates. Replication assays were conducted with three independent biological replicates, with each biological replicates including three technical replicates. Results from all biological replicates, each consisting of technical triplicates, were pooled and presented together in single graphs for analysis. For quantification of schizont and ring-stage parasitemia by air-dried thin smears, a minimum of 10 random fields was counted per sample, totaling 1,000 cells analyzed across three independent biological replicates. The angles between two SPMTs were measured in three independent biological replicates. For each biological replicate, three schizonts per condition were analyzed, with a minimum of 20 pairs of SPMTs per schizont evaluated. Statistical analyses for all experiments were performed using GraphPad Prism software.

## Data availability

All supporting data from this study are available in the main text and the supplementary information files. Parasite strains generated in this study are available upon request to the corresponding author, distribution may require a material transfer agreement (MTA) in accordance with institutional guidelines. Whole genome sequencing data have been deposited in the NCBI Sequence Read Archive (xxx).

## Authors contributions

We acknowledge the support of Ross Tomaino at the Taplin Mass Spectrometry Facility Core. This work was supported by the National Institutes of Health (R01 AI169648 to J.D.D.) and the American Heart Association Postdoctoral Fellowship (25POST1373695 to P.G.).

P.G. was responsible for conceptualization, methodology, formal analysis, investigation, writing—original draft preparation, visualization, and funding acquisition, P.B. performed iterative U-ExM experiments and analysis. I.A. and G.H. generated the PfBBx^HA^ parasite strain and performed IFA on this strain. J.D.D. contributed to conceptualization, data curation, supervision, project administration, funding acquisition, and writing -review and editing.

