## Extended_Data_Figures for "The essential function of the apical polar ring during the blood stage of *Plasmodium falciparum*"

### Extended Data as Supplementary

a

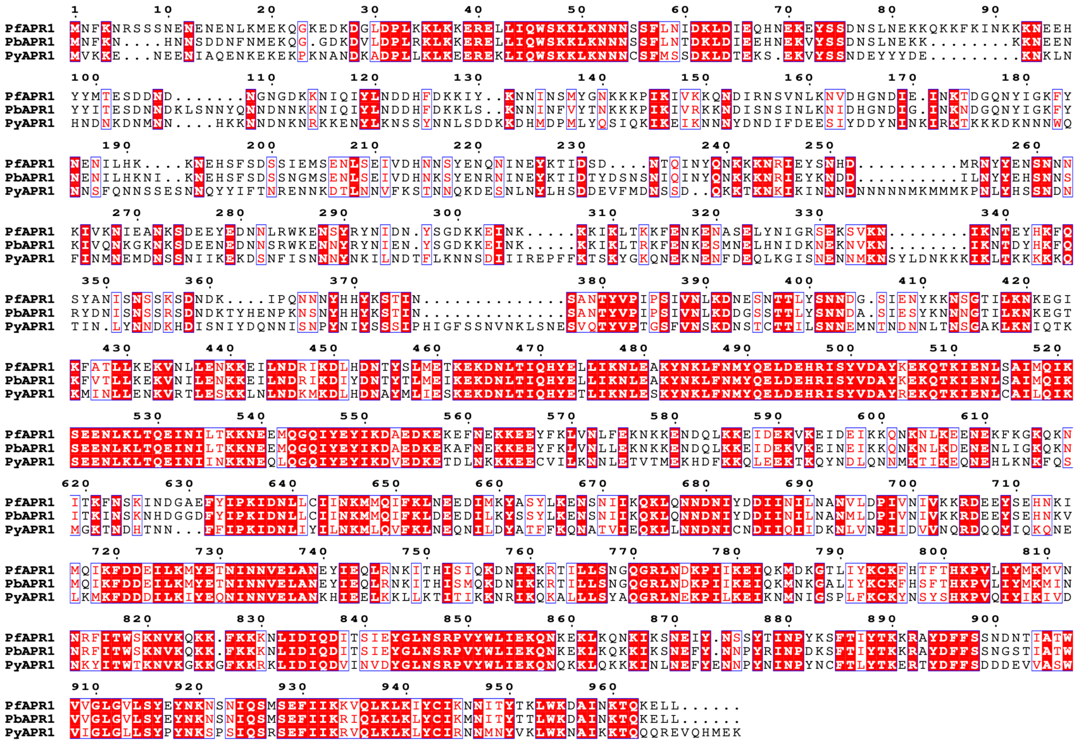

b

| | Sequence Identity | Sequence Similarity |
|---|---|---|
| PbAPR1 | 44.9% | 63.3% |
| PyAPR1 | 45.7% | 62.2% |

**Extended Data Fig. S1: Sequence alignment of PfAPR1 and its orthologs in rodent malaria parasites. a**, Sequence alignment of PfAPR1 (*Plasmodium falciparum*: PF3D7_1141300), PbAPR1 (*P. berghei*: PBANKA_0907700), and PyAPR1 (*P. yoelii*, Py17XNL_000900117) was generated using CLUSTALW in Mega 12 software and visualized with ESPript3. **b,** Sequence coverage and sequence identity between PfAPR1 and its orthologs were determined by BLASTp and EMBOSS Needle analysis.

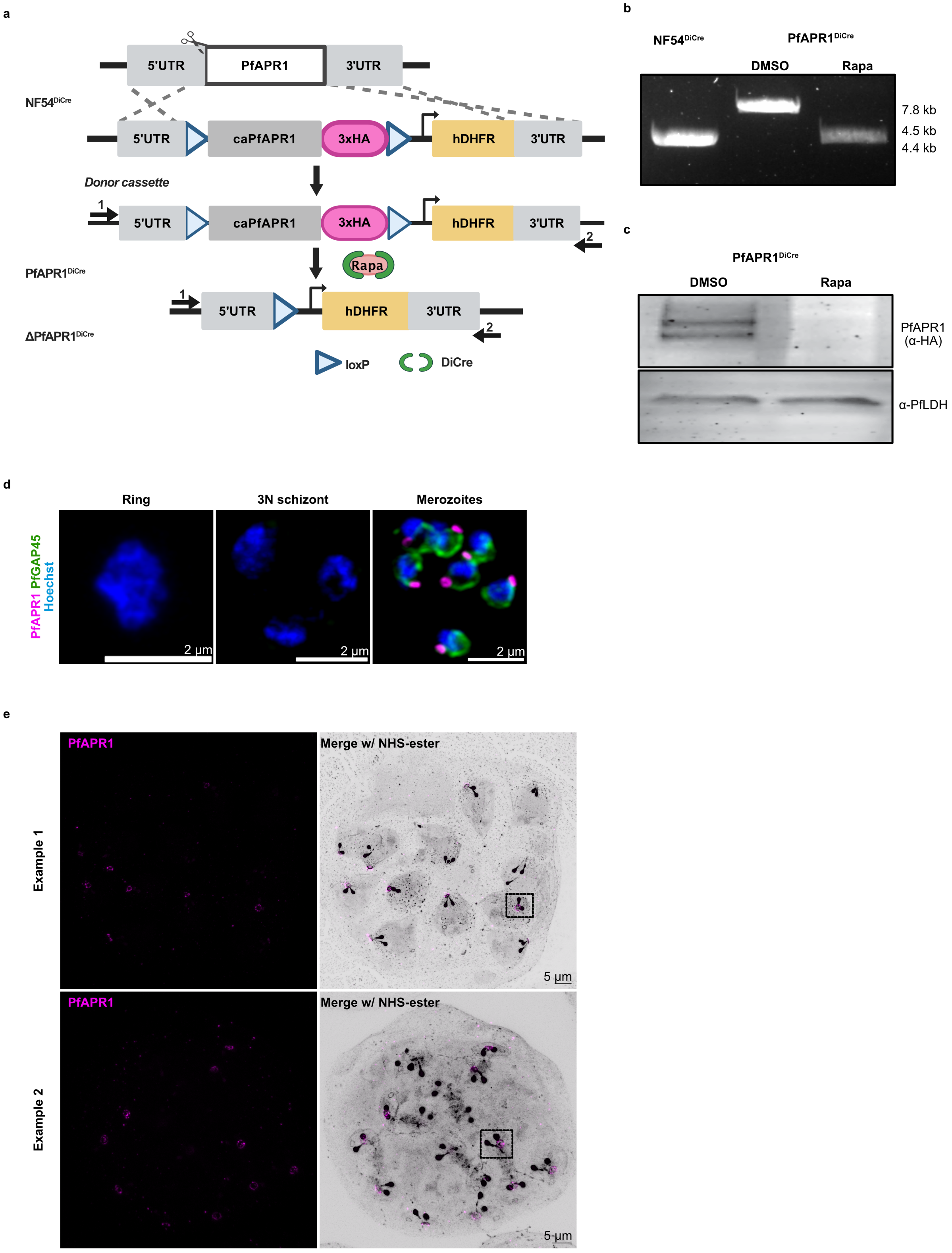
a
5'UTR
PfAPR1
3'UTR
NF54DiCre
5'UTR
caPfAPR1
3xHA
hDHFR
3'UTR
Donor cassette
1
2
PfAPR1DiCre
Rapa
ΔPfAPR1DiCre
loxP
DiCre
b
NF54DiCre
PfAPR1DiCre
DMSO
Rapa
7.8 kb
4.5 kb
4.4 kb
c
PfAPR1DiCre
DMSO
Rapa
PfAPR1
(α-HA)
α-PfLDH
d
Ring
3N schizont
Merozoites
PfAPR1 PfGAP45
Hoechst
2 µm
e
PfAPR1
Merge w/ NHS-ester
Example 1
Example 2
5 µm

**Extended Data Fig. S2: Generation of an inducible knockout (iKO) parasite strain for PfAPR1:** **a,** Schematic representation of the construct and strategy used to generate the PfAPR1$^{DiCre}$ parasite strain. The endogenous PfAPR1 locus in the NF54$^{DiCre}$ strain was replaced with a donor cassette containing a loxP-flanked, codon-altered PfAPR1 fused to a C-terminal 3xhemagglutinin (HA) tag, using CRISPR-Cas9. The human dihydrofolate reductase (hDHFR) serves as a selectable marker. The NF54$^{DiCre}$ strain expresses two halves of the dimerizable Cre recombinase [1]. Addition of rapamycin dimerizes Cre recombinase, resulting in excision of the loxP-flanked PfAPR1 gene. Tightly synchronized ring staged parasites (< 4hpi) are treated with 100 nM of rapamycin (Rapa) for overnight to induce PfAPR1 gene knockout (KO) and DMSO was used as a vehicle control. Excision and loss of PfAPR1 are confirmed by PCR and western blot analysis. **b,** PCR verification of PfAPR1 locus integration and excision in PfAPR1$^{DiCre}$ parasite strain. Expected amplified product sizes are 4.4 kb (native PfAPR1 locus in NF54$^{DiCre}$), 7.8 kb (modified PfAPR1 locus in PfAPR1$^{DiCre}$), and 4.5 kb (rapa-excised PfAPR1 locus in PfAPR1$^{DiCre}$), obtained using primers 1 (oJDD7638) and 2 (oJDD7640). **c,** PfAPR1 expression (~116 kDa) and loss are confirmed by western blot probed with α-HA (3F10, 1:1000). A higher molecular weight band is also detected. α-PfLDH (1:2000) serves as a loading control. **d,** Immunofluorescence assay was used to assess the PfAPR1 expression in parasite stages different from those shown in the main figure (Fig1), specifically in rings, schizonts (3 nuclei stage), and free merozoites using α-HA antibody (1:250, magenta). PfAPR1 expression is detected only in free merozoites (and schizonts, shown in main figures). An α-PfGAP45 (1:500, green) is used as inner membrane complex (IMC) associated protein marker, and Hoechst (blue) stains nuclei. The scale bar represents 2 μm (white bar). **e,** Maximum projection images of full schizont from iU-ExM, showing two examples (boxed region) as presented in Fig. 1d. PfAPR1(α-HA, magenta) and NHS-ester (grey).. Scale bar 5 μm (black bar).

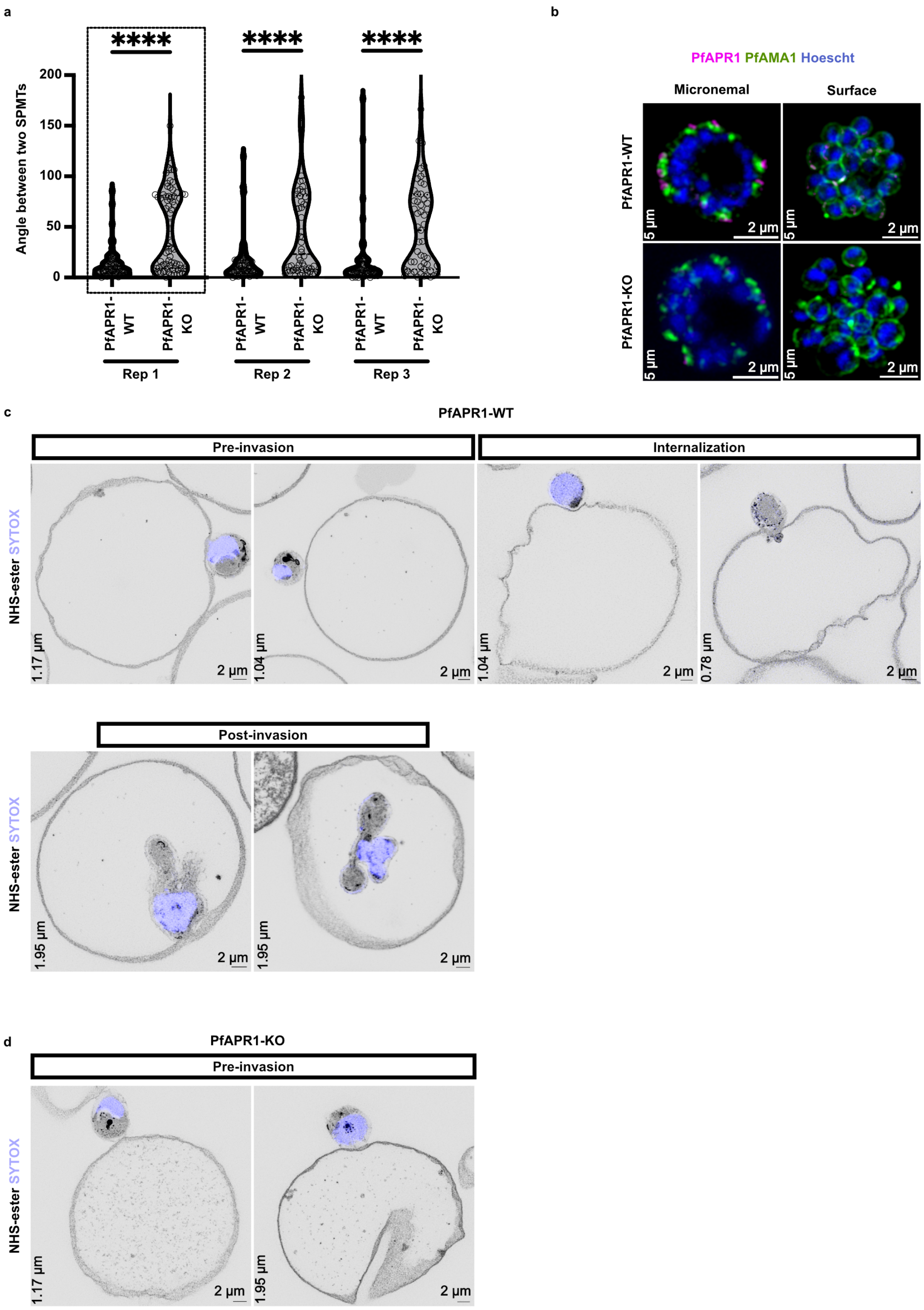
a
b
****
****
****
Angle between two SPMTs
200
150
100
50
0
PfAPR1-WT
PfAPR1-KO
Rep 1
Rep 2
Rep 3
PfAPR1 PfAMA1 Hoescht
Micronemal
Surface
PfAPR1-WT
PfAPR1-KO
5 µm
2 µm
c
PfAPR1-WT
Pre-invasion
Internalization
NHS-ester SYTOX
1.17 µm
1.04 µm
1.04 µm
0.78 µm
2 µm
Post-invasion
1.95 µm
1.95 µm
d
PfAPR1-KO
Pre-invasion
1.17 µm
1.95 µm

**Extended Data Fig. S3: Impact of PfAPR1-KO on SPMT organization and invasion process. a,** ML10 stalled PfAPR1 WT and KO parasites were imaged using U-ExM. The angle between two SPMTs in control and PfAPR1-KO parasites was calculated for each SPMT using the Neuroanatomy plugin (SNT) in FIJI. The graph represents data from three independent biological replicates, the replicate under dotted box is represented in **Fig. 3 c**. For each condition, three schizonts per replicate and at least 20 SPMT pairs per schizont were analyzed. **b,** Early and E64-stalled PfAPR1$^{DiCre}$ parasites were used to assess the translocation of micronemal protein PfAMA1 to the surface of mature merozoites at the end of schizogony in both PfAPR1-WT and KO parasites. IFA demonstrate the PfAMA1 (α-PfAMA1, green) translocation appears normal in both WT and KO parasites. **c** and **d,** Representative U-ExM images depict the invasion process - including pre-invasion, internalization, and post-invasion stages- in PfAPR1-WT and KO parasites. Zoomed views of these images are shown in Fig. 4e. NHS-ester is used as a general protein stain and SYTOX is used to stain nuclei. The images are representative of three independent biological replicates. The scale bar represents 2 μm (white); z-stack depth is shown in the left corner of each image. **d,** All images are representative of a minimum of three independent biological replicates.

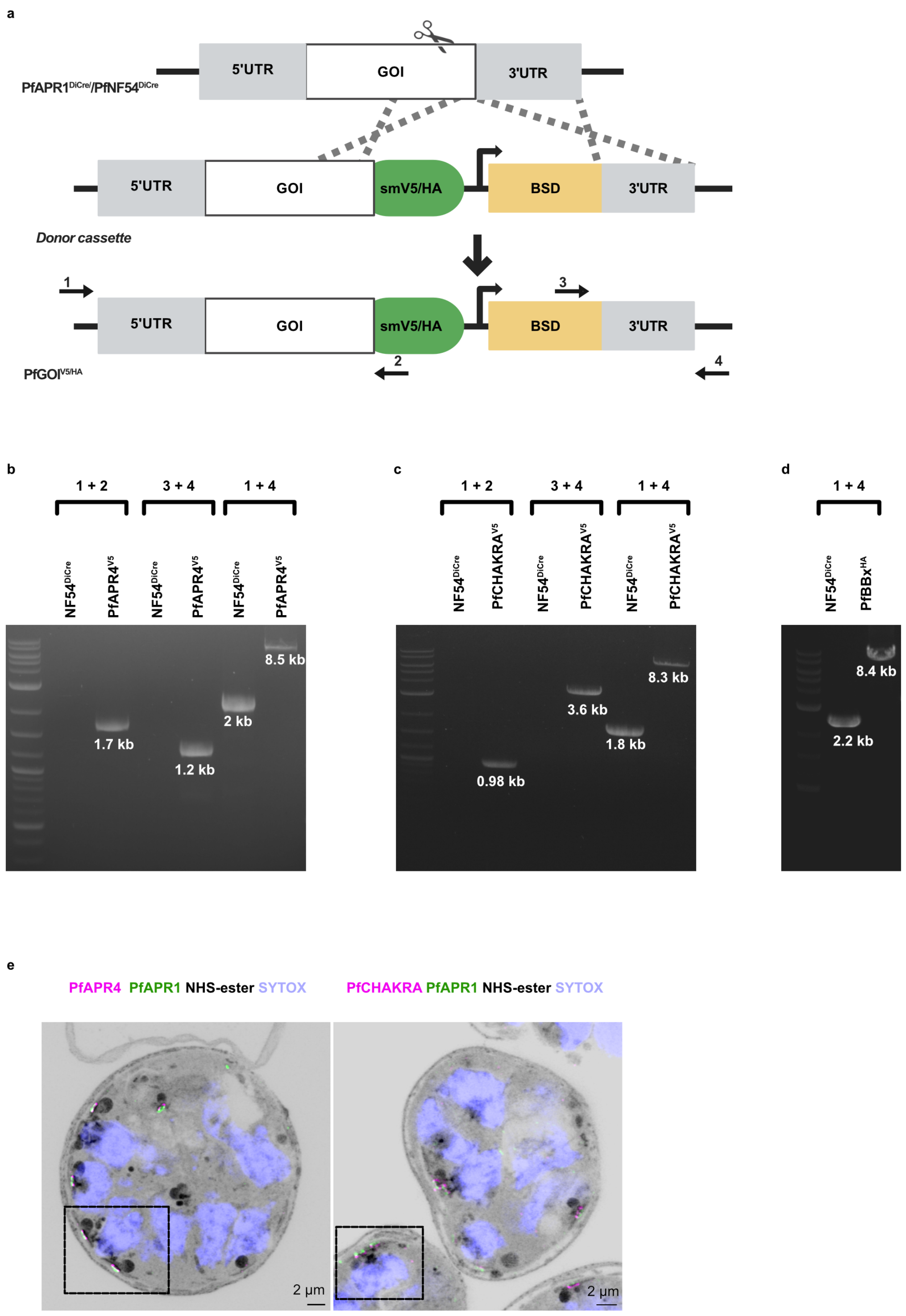
a
5'UTR
GOI
3'UTR
PfAPR1DiCre/PfNF54DiCre
5'UTR
GOI
smV5/HA
BSD
3'UTR
Donor cassette
1
3
5'UTR
GOI
smV5/HA
BSD
3'UTR
2
4
PfGOIV5/HA
b
1 + 2
3 + 4
1 + 4
NF54DiCre
PfAPR4V5
NF54DiCre
PfAPR4V5
NF54DiCre
PfAPR4V5
8.5 kb
2 kb
1.7 kb
1.2 kb
c
1 + 2
3 + 4
1 + 4
NF54DiCre
PfCHAKRAV5
NF54DiCre
PfCHAKRAV5
NF54DiCre
PfCHAKRAV5
8.3 kb
3.6 kb
1.8 kb
0.98 kb
d
1 + 4
NF54DiCre
PfBBxHA
8.4 kb
2.2 kb
e
PfAPR4 PfAPR1 NHS-ester SYTOX
PfCHAKRA PfAPR1 NHS-ester SYTOX
2 μm
2 μm

**Extended Data Fig. S4: Generation of dual transgenic parasites for PfAPR1-interacting candidates. a,** Schematic representation of the construct and strategy used to generate dual transgenic PfGOI$^{V5/HA}$ parasites for three selected candidates: PfAPR4, PfCHAKRA, and PfBBx. The endogenous gene locus in the PfAPR1$^{DiCre}$ strain was replaced with a donor cassette containing GOI fused to a C-terminal spaghetti monster V5 (smV5) or smHA tag, using CRISPR-Cas9. Blasticidin S deaminase (BSD) serves as a selectable marker. PfAPR4 and PfCHAKRA were transfected into PfAPR1$^{DiCre}$ parental background, while PfBBx was integrated into PfNF54$^{DiCre}$. **b,** Integration of the gene in these strains was verified by PCR using primers listed in the **Supplementary Table 2**. **c,** Representative U-ExM images show the localization pattern of PfAPR4 and PfCHAKRA (α-V5, magenta) with respect to PfAPR1 (α-HA, green). The boxed region is shown in Fig. 5. NHS-ester (grey) stains general protein and SYTOX (blue) stains nuclei. The scale bar is 2 μm (white).

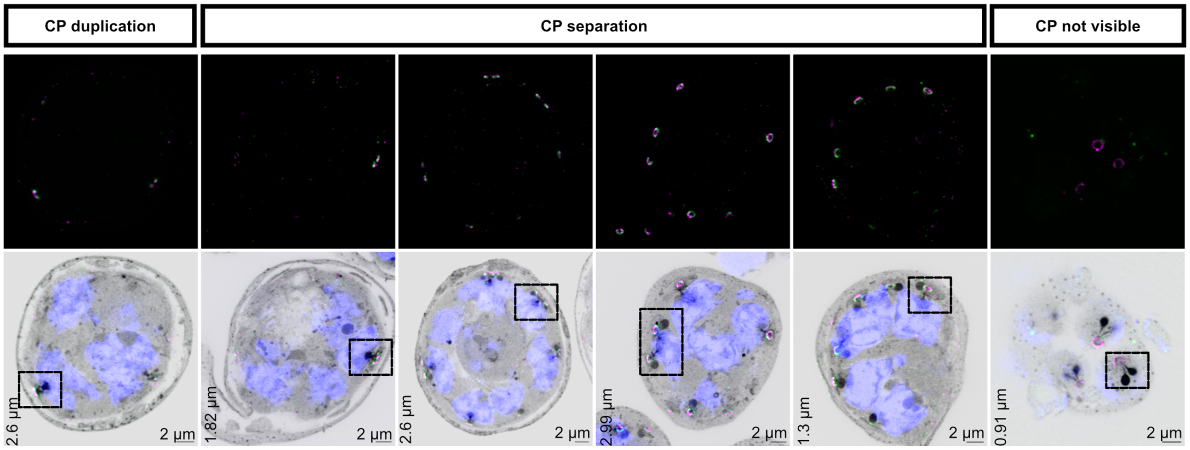


**Extended Data Fig. S5: APR biogenesis using PfAPR1 and PfCHAKRA staining in dual transgenic PfCHAKRA$^{V5}$ parasites**. Representative U-ExM images show APR biogenesis using PfAPR1 (α- HA, magenta) and PfCHAKRA (α-V5, green) The boxed region in these images is shown in Fig. 6.NHS-ester and SYTOX are used as general protein and nuclei stain, respectively. The scale bar is 2 μm (white) and z-stack depth is indicated in the left corner of each image.
