## Supplementary material for "The essential function of the apical polar ring during the blood stage of *Plasmodium falciparum*": Table 2

| Table 2: Oligonucleotides used in the study | | |
|---|---|---|
| **Name** | **Sequence** | **Purpose** |
| **iKO-PfAPR1$^{DiCre}$ parasite strain** | | |
| **To construct pPG35 plasmid containing donor cassette for iKO-PfAPR1$^{DiCre}$ parasite strain** | | |
| oJDD7410 | GTACAGGGTCTCGCCGCTATAATATCC ATTTCAGTCTACTAGAAAGAAAAC | Amplify 5'UTR region of PfAPR1 from Pf3D7 genomic DNA |
| oJDD7411 | GTACAGGGTCTCGTTATTTCGAAAAAC AACAAGAAAAAAAAAAAAAAAAAAAAAA ATATATATATATATATATATATGAATAGA ATATACAACTCAC | |
| oJDD7412 | GTACAGGGTCTCCATAACTTCGTATAG CATACATTATACGAAGTTATATGGTAAA GAAAGAAAACGAGGAAAAC | Amplify codon-optimized PfAPR1 from a geneblock GB85 |
| oJDD7413 | GTACAGGGTCTCCATGGTTTTTCCATG TGTTGAACCTCTCTC | |
| oJDD7414 | GTACAGGGTCTCGCCATGGGGGGGCG GTGGTTCCGGTG | Amplify region containing linker, 3HA_Loxp. nLuc, hDHFR from plasmid pPG31 |
| oJDD7415 | GTACAGGGTCTCGAAGCTAGATTTAAT AAATATGTTCTTATATATAATGAG | |
| oJDD7416 | GTACAGGGTCTCGGCTTCTCGAGGCG CCAATCATATTATCAACACATTTG | Amplify 3' UTR of PfAPR1 from Pf3D7 genomic DNA |
| oJDD7417 | GTACAGGGTCTCGTCAGCTCGAGGAA TTCTACATCTTTACGGATTATATTATTC ATACGGC | |
| oJDD7418 | GTACAGGGTCTCGCTGATGCGGTATTT TCTCC | Amplify pGEM backbone from plasmid pPG25 |
| oJDD7419 | GTACAGGGTCTCGGCGGCCGCACGCG TCAGAGTATTCTATAGTG | |
| **For guide RNA plasmids** | | |
| oJDD7420 | TATTGAAAGAGAAAAATTAATCCAG | pPG36 for PfAPR1$^{DiCre}$ parasite strain |
| oJDD7421 | AAACCTGGATTAATTTTTCTCTTTC | |
| **For Integration check** | | |
| oJDD7638 | ATTATCCGTGAAATATATTTC | Whole locus integration check (7757 bp) |
| oJDD7640 | GAAATATTTTGAGCAGGAAAGG | |
| **Geneblock GB85** | | |

| | |
|---|---|
| GB85 | atggtaaagaaagaaaacgaggaaaacatagcacaagaaaacaaagagaaggagaagcctaaaaatgctaat<br>gacaaggccgatccacttttaaaattgaaggaggagcgtgagaagttaatacagtggtcaaaaaaattaaaaataa<br>caactgctcattcatgtcatctgataagttggacaccgagaaaagtgagaaggtttattcttctaatgacgagtattactac<br>gacgagaagaacaaattaaaccacaatgacaacaaagataacatgaacaaccataagaagaataatgataacaa<br>aaacaggaagaaagaaaactacttgaagaatagttcttacaataacctttcagatgataagaaagatcacatggacc<br>ctatgttataccaatcaattcaaaaaattaaagagattaaaaataataattatgacaacgacattttcgatgaagaatca<br>atatacgatgactataatattaataaaatacgtaagactaagaaaaaggataaaaacaacaattggcagaataactc<br>attccagaataacagttctgagtctaataaccaatactacatattcacaaaccgtgagaataataaagacacattgaac<br>aatgttttcaaatctacaaataatcagaaagatgaaagtaaccttaactatttacacagtgatgacgaggtattcatggat<br>aacagttcagatcagaaaaaaactaaaaataaaataaaaattaacaacaacgataacaacaacaacaatatgaaa<br>atgatgatgaagcctaacttgtaccactcatctaatgataattttattaatatgaacgaaatggataactcatctaatataat<br>aaaagagaaggactcaaatttcatttcaaacaacaattacaataaaatattaaatgacacatttcttaaaaataactcag<br>atattataatacgtgagcctttttttaaaacttctaagtacggaaagcaaaatgagaaaaacgaaaactttgatgagcaa<br>cttaaaggtatatctaatgaaaacaatatgaaaaattcttacttggataataagaaaaaaattaagttgacaaaaaaaa<br>aaaagaaacagaccattaacttatacaacaacgataagcacgacataagtaatatatatgatcagaataatatttcaa<br>atccatacaatatttattcttcttcaattccacatattggtttttcttctaacgttaacaaattatctaatgaatcagtacaaacct<br>acgtaccaacaggaagtttcgttaattctaaagataacagtacctgtaccactattctttcaaataatgagatgaatacaa<br>acgataacaacttgaccaacagtggagcaaaattaaaaaacatacaaacaaaaaagatgattaacttacttgagaa<br>caaagttaggaccttggaatctaaaaaattaaacttaaacgacaagatgaaggatttgcatgataatgcctacatgttga<br>tagagtcaaaggaaaaagacaaccttactattcaacattatgaaacacttataaagaacttggaatctaagtataataa<br>attgtttaacatgtaccaggaacttgacgagcatcgtatatcatacgtggatgcctatcgtgaaaaacagacaaagata<br>gaaaacttatgtgccatattacagataaaatctgaggagaacttaaaattgactcaggagataaatataataaataaga<br>aaaatgaacaattgcagggtcagatttatgagtacattaaggacgtggaagacaaggaaacagacttgaacaagaa<br>aaaagaagagtgcgttattcttaaaaataacttggagacagtaacaatggaaaagcacgacttcaaaaagcagttag<br>aggaaaagaccaagcagtacaacgatttacagaacaacatgaagaccattaaagagcaaaatgaacaccttaag<br>aacaagttccaatcaatgggaaaaaccaacgaccatactaacaatttctttataccaaagatagacaatttaatttatatt<br>ttaaataagatgcttcaggtgtttaaattgaacgagcagaacattttggattatgccacattcttcaaacaaaatgcaaca<br>gttattgaacagaaacttttgaataacgataatatttgtaatgatattatacagataatagataagaacttagtgaatccaat<br>tatagatgttgtaaaccagagggatcagcagtatattcagaagcagaatgaattgaaaatgaagttcgacgatgacatt<br>cttaagatatatgaacaaaacataaacaacgtggagttagccaacaaacatattgaagagcttaagaaattacttaaa<br>accataactataaaaaaaaataggattaagcaaaaagcattgcttttatcttatgctcaaggaaggttaaatgagaagc<br>caatattaaaagaaataaaaaacatgaacataggaagtcctttgtttaagtgtaaatacaatagttactctcacaagcca<br>gttcagatatatataaagattgtagataacaagtatattacctggacaaaaaatgttaaaggtaaaaagggtttcaaaaa<br>gaggaagttaatagacattcaagacgtgattaatgtagactacggattaaacagtcgtccagtgtattggttgattgaaa<br>agcaaaatcaaaaaaaacttcagaaaaaaaagataaatcttaacgagttttatgagaataatccatataacataaacc<br>catacaattgtttcacattgtatactaaggagaggacatacgatttcttcagtgacgatgatgaggtagtggcatcatgggt<br>tattggattaggtcttttgtcatacccttacaataaatctccttctattcaaagtaggagtgagtttattataaagagggtgcaa<br>ttgaaacttaagttatactgcattaggaacaacatgaattacgtaaagttatggaaaaacgctataaagaagactcaac<br>agcagagagaggttcaacacatggaaaaataa |

**PfAPR4$^{V5}$ parasite strain**

**To construct pPG61 plasmid containing donor cassette for PfAPR4$^{V5}$ parasite strain**

| | | |
|---|---|---|
| oJDD8357 | GTACAGGGTCTCGCCGCGAGAATAAT<br>GTAAAAATGGATAAG | Amplify 5' homology region (HR) region from Pf3D7 genomic DNA |
| oJDD8358 | ATAAATAAATTTGTTGATGTTCGATG | |
| oJDD8463 | CATCGAACATCAACAAATTTATTTATAC<br>AAATAACGAGACAGTTCAG | Amplify a codon-optimized PfAPR4 gene from geneblock GB100 |

| | | |
|---|---|---|
| oJDD8464 | GGAAAACTTTGAATTATTACAAGAAAATATATTGAAAGAGGACGAACAAG | |
| oJDD8331 | GTACAGGGTCTCGGGGGGCGGTGGTTCCGGTGGTGGTGGTTCTCCATGGATGGGAAAACCTATAC | Amplify smV5 from pPG03 |
| oJDD8332 | GTACAGGGTCTCGTAAGTATTTGATGAATTAACTACACTTAAAATAATAC | |
| oJDD8333 | GTACAGGGTCTCGCTTAAGCGGAAAGGGGCCATTGG | Amplify BSD from pPB69 |
| oJDD8334 | GTACAGGGTCTCGGAAATTGAAGGAAAAACCATCATTTGTG | |
| oJDD8361 | GTACAGGGTCTCGTTTCGATATGTTTTTTTTTTTTTTTTTTTTTTTTTTAATTGATTATTCATTTTG | Amplify a 3’ HR region from Pf3D7 genomic DNA |
| oJDD8362 | GTACAGGGTCTCGATTCGTATTTATATTTACCTCTCCCC | |
| oJDD8337 | GTACAGGGTCTCGGAATTCCTCGAGCTGATGCGGTATTTTCTCC | Amplify the pGEM backbone from pPG25 |
| oJDD8338 | GTACAGGGTCTCGGCGGCCGCACGCGTCAGAGTATTCTATAGTG | |
| **For guide RNA plasmids (pPG62, pPG63, and pPG64)** | | |
| oJDD8403 | TATTGAACAATGAAACGGTACAACA | pPG62 |
| oJDD8404 | AAACTGTTGTACCGTTTCATTGTTC | |
| oJDD8405 | TATTGACAATCAGATGACATGAATG | pPG63 |
| oJDD8406 | AAACCATTCATGTCATCTGATTGTC | |
| oJDD8407 | TATTGATATAAGAGTTATCGCAAGG | pPG64 |
| oJDD8408 | AAACCCTTGCGATAACTCTTATATC | |
| **For Integration check** | | |
| oJDD8644 | GGGTTAAAGAGGAAACGGATGC | 5' integration check (1695 bp) |
| oJDD5365 | CTTCCACGTTGTGCCGAATTTT | |
| oJDD0832 | CTTTACACACATAAAATGGCTAGTATG | 3' integration check (1283 bp) |
| oJDD8645 | TCCAAAAGGGATAATATTCATACAGTAC | |
| oJDD8644 | GGGTTAAAGAGGAAACGGATGC | Modified region check (8516 bp) |
| oJDD8645 | TCCAAAAGGGATAATATTCATACAGTAC | |
| **Geneblock GB100** | | |

| | | |
|---|---|---|
| GB100 | cctttgaagattataaatggtaagaacaaacttataaaggacttgaagaaaattcaggagcaggtagagaggaaaat acgtaaatacaaaatacagatggaccaggagaacaagaagccacctcctagtaagaataagataaacatgaagtc aattaacttagatatagacgacgaccagaacgttgactcacagggtatagtggactatgtgcttaaccaaataggtaat aagaagatgggacaaacaaataacgagacagttcagcaagacgtattcaacagtgctgacattaaggagttgtacct taaagttcaagttccaccatgtgacaatagttacatgtcatctagtgttcagtctgacgatatgaacgaagaggaatatga gaagtatgatttaattaaccacgcccaaaaaaagaaacaaaacggaatacctgtaaagattaagtacgttcgtgaca cattggacgacgagaatagtaagtatgtgatagaaattcctaaaaaagaccttaagggacacgaaaaaaaggatca cataaaagctgtattatggggtaattcaaaaaagggtcttgaaattaagcttccaggtactcataacttcattcctcagatg aagataaaagagcaaattttaattcagagaggattattgaaaataaaagagcaacagcaagtttatttggaattacttaa aaaaaacgacataaagcaacataaggagaatatacacaacattatt | |
| **PfCHAKRA$^{V5}$ parasite strain** | | |
| **To construct pPG72 plasmid containing donor cassette for PfCHAKRA$^{V5}$ parasite strain** | | |
| oJDD8375 | GTACAGGGTCTCGCCGCCCACATATG CTGATAAGGTTTTAG | Amplify a 5’ HR region from Pf3D7 genomic DNA |
| oJDD8376 | ATTATTTATCTGTTTATTCAAATCTG | |
| oJDD8466 | CAGATTTGAATAAACAGATAAATAATCC TTTAAGTCACTTGAACTTAAATG | Amplify a codon-optimized region from geneblock GB102 |
| oJDD8378 | GTACAGGGTCTCGCCCCACCAAGTATT CCCTCTTTTTTAAGG | |
| oJDD8331 | GTACAGGGTCTCGGGGGGCGGTGGTT CCGGTGGTGGTGGTTCTCCATGGATG GGAAAACCTATAC | Amplify smV5 from pPG03 |
| oJDD8332 | GTACAGGGTCTCGTAAGTATTTGATGA ATTAACTACACTTAAAATAATAC | |
| oJDD8333 | GTACAGGGTCTCGCTTAAGCGGAAAG GGGCCATTGG | Amplify BSD from pPB69 |
| oJDD8334 | GTACAGGGTCTCGGAAATTGAAGGAAA AACCATCATTTGTG | |
| oJDD8379 | GTACAGGGTCTCGTTTCAAATGCAATA ATCAAATTTGAGACATTATAC | Amplify 3’ HR from Pf3D7 genomic DNA |
| oJDD8380 | GTACAGGGTCTCGATTCGTACAGCAAA TTTAATTCGTTACCAG | |
| oJDD8337 | GTACAGGGTCTCGGAATTCCTCGAGCT GATGCGGTATTTTCTCC | Amplify the pGEM backbone from pPG25 |
| oJDD8338 | GTACAGGGTCTCGGCGGCCGCACGCG TCAGAGTATTCTATAGTG | |
| **For guide RNA plasmids (pPG73 and pPG74)** | | |
| oJDD8419 | TATTGTTAAGATTTAAGTGACTGAG | pPG73 |
| oJDD8420 | AAACCTCAGTCACTTAAATCTTAAC | |
| oJDD8421 | TATTGTCATTCAATAAAAGAGACCA | pPG74 |
| oJDD8422 | AAACTGGTCTCTTTTATTGAATGAC | |
| **For Integration check** | | |

| | | |
|---|---|---|
| oJDD8566 | CAATGAATATAGAAGAAGGTGCAAG | 5' integration check (980 bp) |
| oJDD2933 | CTGCTGCTGAGTACTATCAAGTC | |
| oJDD8539 | CGTAATAACGCTAAAAGTTTTAGATGTGCTTTACTAAGTC | 3' integration check (3589 bp) |
| oJDD8567 | CAATATTACCTTTCGTAACATTTGTTC | |
| oJDD8566 | CAATGAATATAGAAGAAGGTGCAAG | Modified regions check (8333 bp) |
| oJDD8567 | CAATATTACCTTTCGTAACATTTGTTC | |
| **Geneblock GB102** | | |
| GB102 | ccatcttgcagtccttctggttgcgcttcaaggaataaagtaactcatgtgcagaatgtgaggagaaactcatatataaatccacattcacactcaatgcacgagtgcccaaaaaaagtgcaaattattgcacgtcctatgcacgatcagaatcttcgtacctattctctttgtcgtatattgaaagaggacgaacaagggcttccaataaacaatacaccatttttcataaagggaaagttgctttgattccaaaagacataaatacatttttcgacgatgaaaaaaagggagaaagtacaggtaagaaagactactacgccagacaggtagtacagatggccacacaggtaaatactcctacacctgtgaaatacgttccaaagtttacaaaagctaaggtgaaacctaaggtgcctcttttattaaaaaatataacaaaggctcctttaagtcacttgaacttaaatgatttttcaaacattcattatcacgacttcaatcttcagaacaccaccaacaacaagcagttccgtaaatacagtactaacggatacaataacatttacaatgagcttaacaagaacgttgctccagatattttctcttttaacaagagggatcagggtaattattttaagtcagagacaagaaagagtttgagtagttttagggaagcccttaaaaaagagggaatacttggt | |
| **PfBBx$^{HA}$ parasites** | | |
| **To construct pIA39 plasmid containing donor cassette for PfBBx$^{HA}$ parasites** | | |
| oJDD8875 | GTACAGGGTCTCGAGGCCTTGCAATTTGTTCAAACGAATC | Amplify a 5' HR region amplified from Pf3D7 genomic DNA |
| oJDD8876 | GTACAGGGTCTCGGGTATGCTTGCAAGTACTTTAAATATGAATTAATTTGTGATGCATAAAAATCCATATAG | |
| oJDD8877 | GTACAGGGTCTCGTACCTTAATGAGTCAAATATGTTGGATAGTAAAGGGGGCGGTGGTTCC | Amplify a smHA and BSD from pPB69 |
| oJDD8878 | GTACAGGGTCTCGGAAATTGAAGGAAAAACCATCATTTGTG | |
| oJDD8879 | GTACAGGGTCTCGTTTCTCAAAAAATGTATACATACACTTCATAC | Amplify a 3' HR region from Pf3D7 genomic DNA |
| oJDD8880 | GTACAGGGTCTCGATGGCTTCGGTGTTCATTTTAAATTTTC | |
| oJDD8818 | GTACAGGGTCTCGCCATGGCTGATGCGGTATTTTCTCCTTACG | Amplify the pGEM backbone from pPB69 |
| oJDD8881 | GTACAGGGTCTCGGCCTGAGTATTCTATAGTGTCACCTAAATAGCTTG | |
| **For guide RNA plasmids (pIA44 and pIA45)** | | |
| oJDD8578 | TATTGTAGTTATCTTAAATATCTAC | pIA44 for PfBBx$^{HA}$ parasites |
| oJDD8579 | AAACGTAGATATTTAAGATAACTAC | |
| oJDD8580 | TATTGTACAAATATTGGATAACCAT | pIA45 for PfBBx$^{HA}$ parasites |

| | | |
|---|---|---|
| oJDD8581 | AAACATGGTTATCCAATATTTGTAC | |
| **For Integration check** | | |
| oJDD8887 | GGTGATGATGACAATGTATGTAG | Whole locus integration check |
| oJDD8889 | TCATCTTCAGCATCATTTGATG | |
