## Supplementary material for "The essential function of the apical polar ring during the blood stage of *Plasmodium falciparum*": Table 3

### Table 3: Summary of all antibodies and stains used in this study

#### Primary Antibody

| Name | Ab species | Source (CatLog number) | Dilutions used | References |
|---|---|---|---|---|
| α-PfLDH (clone 19G7+6C9) | Mouse | Mouse anti-LDH (pan reactive for plasmodium species) from Mike Makler at Flow Inc | Western Blot control for parasite lysate (1:2000) | |
| α-HA (3F10) | Rat | Sigma 11867423001 | Western Blot (1:1000), IFA (1:250), and U-ExM (1:125) | |
| α-PfGAP45 | Rabbit | gifted by Julian Rayner | IMC marker; IFA (1:5000), U-ExM (1:2500) | Jones M.L. et al, Mol and Biochem Parasitology, 2009 |
| α-PfAMA1 (1F9) | Mouse | gifted by Robin Anders | Apical complex and microneme marker; IFA (1:200) | Coley, et al. Protein Eng, 14:691-698 2001. |
| α-Histone H3 | Rabbit | Abcam (1791) | Western blot control for parasite lysate; IFA (1:2500) | |
| alpha-tubulin (12G10) | Mouse | DSHB (AB_1157911) | Labeling SPMTs; IFA (1:1000) and U-ExM (1:500) | |
| α-V5 (clone SV5-pk1) | Mouse | Biorad Serotech MCA1360 | IFA (1:500) and U-ExM (1:250) | |

#### Secondary Antibody

| Name | Ab species | Source (CatLog number) | Dilutions used | References |
|---|---|---|---|---|
| Alexa 555 goat anti-rabbit | Goat | Thermofisher (A21429) | IFA (1:1000); U-ExM (1:500) | |
| Alexa 488 goat anti-mouse IgG2a | Goat | Thermofisher (A21131) | IFA (1:1000); U-ExM (1:500) | |
| Alexa 488 goat anti-rat | Goat | Thermofisher (A11006) | IFA (1:1000); U-ExM (1:500) | |
| Alexa 555 goat anti-mouse IgG2a | Goat | Thermofisher (A21137) | IFA (1:1000); U-ExM (1:500) | |
| Alexa 555 goat anti-mouse IgG2b | Goat | Thermofisher (A21147) | IFA (1:1000); U-ExM (1:500) | |
| Alexa 488 goat anti-rabbit | Goat | Thermofisher (A11034) | IFA (1:1000); U-ExM (1:500) | |
| Alexa 555 goat anti-mouse | Goat | Thermofisher (A21424) | IFA (1:1000); U-ExM (1:500) | |

| | | | |
|---|---|---|---|
| Alexa 488 goat anti-mouse IgG1 | Goat | Thermofisher (A21121) | IFA (1:1000); U-ExM (1:500) |

| Stains/Dyes | | |
|---|---|---|
| **Name** | **Source (catalog number)** | **Purpose and dilutions used** |
| Hoescht 33342 | ThermoFisher (H3570) | Stains nuclei, IFA (1:2500) |
| Alexa Fluor 405 carboxylic acid, succinimidyl ester (NHS-Ester) | Life Tech (A30000) | General protein stain in U-ExM, U-ExM (10 ug/mL) |
| Wheat Germ Agglutinin ,WGA CF®640R | Biotium (29026) | Stains RBC membrane (10ug/ml) |
| SYTOX (Deep Red Fluorescent Nucleic Acid Stain for Fixed cells) | Invitrogen (S11381) | Stains nuclei in U-ExM (1 μM) |
